# Size Matters: A Mechanistic Model of Nanoparticle Curvature Effects on Amyloid Fibril Formation

**DOI:** 10.1101/2021.07.01.450782

**Authors:** Torsten John, Juliane Adler, Christian Elsner, Johannes Petzold, Martin Krueger, Lisandra L. Martin, Daniel Huster, Herre Jelger Risselada, Bernd Abel

## Abstract

The aggregation of peptides into amyloid fibrils is linked to ageing-related diseases, such as Alzheimer’s disease and type 2 diabetes. Interfaces, particularly those with large nanostructured surface areas, can affect the kinetics of peptide aggregation, ranging from a complete inhibition to strong acceleration. While a number of physiochemical parameters determine interface effects, we here focus on the role of nanoparticle curvature for the aggregation of the amyloidogenic peptides Aβ_40_, NNFGAIL, GNNQQNY and VQIYVK. Nanoparticles (NPs) provided a surface for peptide monomers to adsorb, enabling the nucleation into oligomers and fibril formation. High surface curvature, however, destabilized prefibrillar structures, providing an explanation for inhibitory effects on fibril growth. Thioflavin T (ThT) fluorescence assays as well as dynamic light scattering (DLS), atomic force microscopy (AFM) and electron microscopy experiments revealed NP size-dependent effects on amyloid fibril formation, with differences between the peptides. While 5 nm gold NPs (AuNP-5) retarded or inhibited the aggregation of most peptides, larger 20 nm gold NPs (AuNP-20) tended to accelerate peptide aggregation. Molecular dynamics (MD) studies demonstrated that NPs’ ability to catalyze or inhibit oligomer formation was influenced by the oligomer stability at curved interfaces which was lower at more highly curved surfaces. Differences in the NP effects for the peptides resulted from the peptide properties (size, aggregation propensity) and concomitant surface binding affinities. The results can be applied to the design of future nanostructured materials for defined applications.

## Introduction

Nanostructured materials are abundant in nature, for instance as vesicles and biological membranes in organisms,^1^ or as nanoparticles in air pollutants,^2^ cosmetics^3^ and medical products.^4^ Their high surface-to-volume ratio implies a significant impact of even small amounts upon various biological processes.^5–8^ Of particular interest is the role of nanoparticles (NPs) for peptide aggregation into amyloid fibrils.^5,9–11^ Amyloid fibrils are formed *via* a nucleation-polymerization mechanism,^12,13^ and some intermediates are connected with neurodegenerative diseases, such as Alzheimer’s disease, type 2 diabetes, or prion diseases.^14–16^ A comprehensive overview on the impact of nanoparticles on amyloid peptide aggregation has recently been published.^5^

Earlier studies reported contradictory effects of NPs on amyloid fibril formation ranging from a complete inhibition to strong acceleration of the process (Figure 1).^5,17–20^ These paradoxical effects result from the diversity of NPs which are characterized by varying physicochemical properties, determined by their material,^9^ surface chemistry,^21–23^ morphology and size.^24–30^ NP size effects on amyloid peptide aggregation have been the subject of intense research.^18,25–27,29,31,32^ Experimental studies identified an impact on the immediate peptide layer formed around NPs, the corona,^31^ and particularly the peptide’s secondary structure^18,26^ and β-sheet content.^27^ Gao *et al*. reported an acceleration of Aβ_40_ fibrillation by larger NPs whereas smaller NPs and nanoclusters retarded or inhibited amyloid fibril formation.^29^ Theoretical studies using coarse-grained peptide models described an influence of NP curvature on peptide adsorption energies and densities.^24,25,33^ The adsorption energies increased with increasing NP size due to van der Waals attraction and increasingly exposed surfaces.^24^ While these studies provided insights into peptide adsorption affinities and peptide conformations at NPs of varying size, a comprehensive model of how the complex processes of amyloid fibril formation are influenced by NPs is missing.

**Figure 1.**
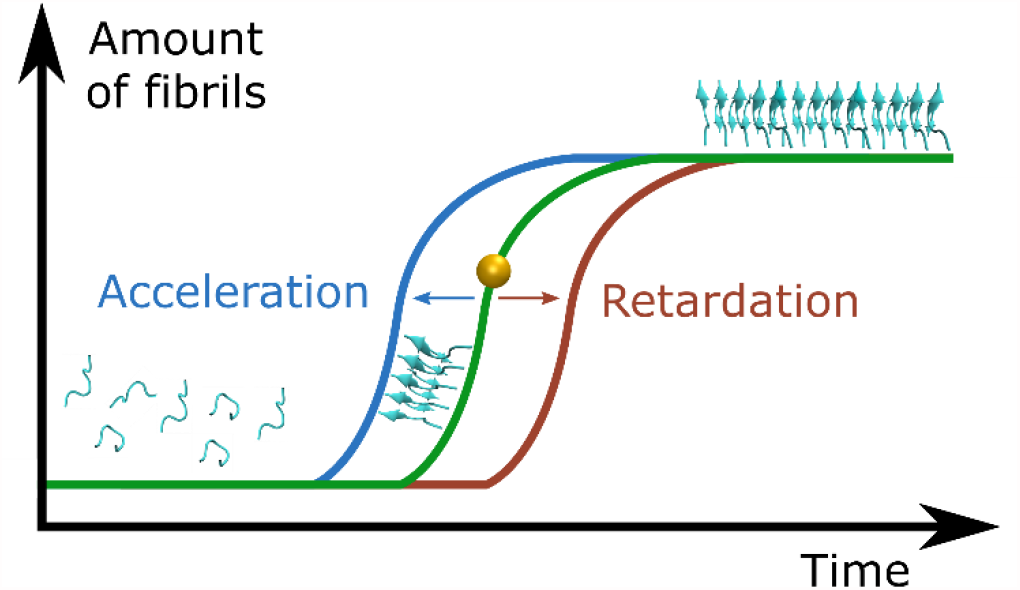
Schematic representation of the contradictory influence of NPs on the kinetics of amyloid fibril formation. Figure adapted from John *et al*.^5^

In addition to NP size, the impact of NPs on amyloid fibril formation also depends on the peptide itself, such as its intrinsic aggregation propensity and fibril flexibility,^34,35^ and the physicochemical environment, such as solution ionic strength, pH and temperature.^36,37^ These properties can be tuned to better determine the potential impact of NPs on disease development or to design NPs with inhibitory effects on amyloid formation.^4,5^ Previous studies found that higher ionic strengths or particular buffer conditions induced or accelerated peptide aggregation due to changes in the strength of electrostatic interactions.^36,38,39^ Due to the influence of all those parameters, it is not unexpected that contradictory effects of NPs on amyloid peptide aggregation have been reported.

In this work, we used experimental techniques in conjunction with molecular dynamics (MD) simulations to demonstrate that NP size does differentially affect fibril growth for several amyloidogenic peptide motifs (amyloid beta peptide: Aβ_40_; human islet amyloid polypeptide: hIAPP, NNFGAIL; prion protein: SUP35, GNNQQNY; tau protein: VQIVYK).^40,41^ Thioflavin T (ThT) fluorescence assays, atomic force microcopy (AFM), dynamic light scattering (DLS) and electron microscopy experiments provided kinetic and structural information on the size-dependent NP influence on amyloid fibril formation, while keeping the physicochemical environment constant. Small 5 nm gold NPs (AuNP-5) retarded or even inhibited the aggregation of most peptides, while larger 20 nm gold NPs (AuNP-20) had either no influence or accelerated peptide aggregation. The curvature effect of NPs on amyloid fibril formation was computationally modelled *via* external potentials that forced fibrillar oligomers on a curved surface. We identified that larger NPs provided an attractive surface for seed formation, while smaller NPs may have interfered with the structure of larger oligomers and fibrils, inhibiting amyloid peptide aggregation. The differential influence of NP size on the aggregation of the peptide motifs studied has been attributed to differences in the peptides’ size and flexibility, aggregation propensity and concomitant surface binding affinity, in a given environment.

## RESULTS AND DISCUSSION

To explore the impact of NP size on amyloid peptide aggregation, gold NPs (AuNPs, citrate-stabilized) of varying diameters (5 nm and 20 nm) were added to buffered peptide solutions. The size and zeta-potential of the NPs (AuNP-5 and AuNP-20) were validated using UV-vis absorption spectroscopy (AuNP-5: 5 nm; AuNP-20: 20 nm) and dynamic light scattering (DLS, AuNP-5: 7 nm (±2), ζ = −41 mV (±10); AuNP-20: 17 nm (±4), ζ = −42 mV (±47)) (Table S1 and Figure S1 for NPs’ characterization). Peptide solutions were buffered in HEPES to ensure a consistent pH, while keeping the citrate-coated AuNPs stable (Figures S2 and S3 and Table S2). This is a critical step to separate surface effects from potential pH changes upon addition of NP solution. Amyloid-forming peptides of varying size (six to 40 amino acids), sequence and overall net charge were studied (Table 1) under consistent conditions (HEPES buffer, 100 mM, pH 7.4, 37°C) to identify general patterns in the NP effects. The peptide concentrations were adjusted so that peptide aggregation could be probed within a reasonable time frame, which differed between the peptides due to their differences in intrinsic aggregation propensity (Figure S4). At 1 mg/mL, only Aβ_40_ aggregated under the study conditions (Figure S5).

**Table 1.**
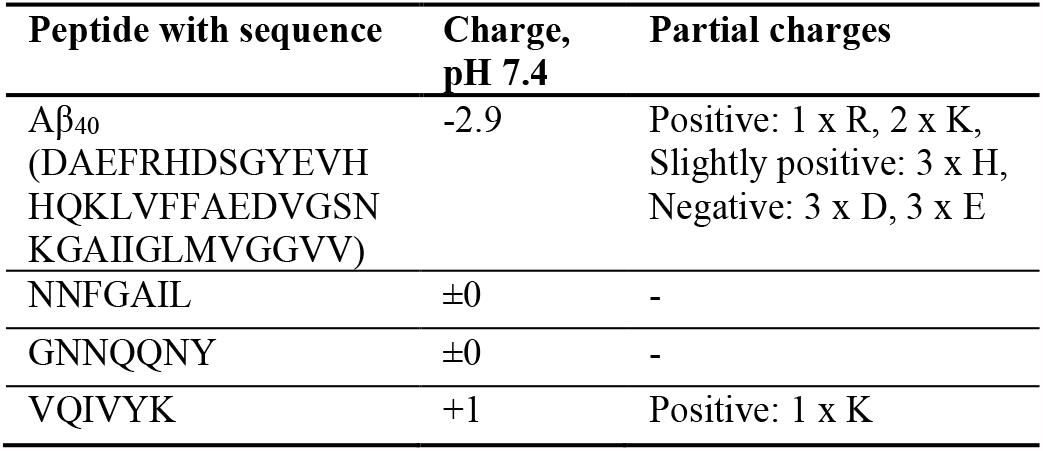
Overview of peptides studied with primary sequence and net charge at pH 7.4 (determined using online calculator^42^).

The kinetics of amyloid peptide aggregation in the presence and absence of NPs was primarily probed using thioflavin T (ThT) fluorescence assays (Figure 2). The ThT dye is commonly used to follow the aggregation kinetics of peptides as it changes its fluorescence properties upon binding to β-sheet rich amyloid fibrils.^43,44^ AuNPs of 5 nm and 20 nm in diameter were added at constant mass (20 µg/mL) and constant overall surface area (3.1 cm^2^/mL). This enabled consideration of the larger surface area at the same mass concentration of the smaller NPs and thus excluding the possibility that NP size-dependent effects would solely result from differences in NP surface area. The smaller NPs (AuNP-5) retarded or even inhibited the aggregation of Aβ_40_, GNNQQNY and VQIVYK aggregation. While the effects were more significant at higher surface area for Aβ_40_ and GNNQQNY (Figure 2 a, b, e), a complete inhibition of the aggregation process was observed for VQIVYK at both 3.1 cm^2^/mL and 12.4 cm^2^/mL AuNP-5 (Figure 2 f). Thus, while the surface area and amount of NPs matter, a size-dependent effect of NPs on peptide aggregation was observed. When comparing the different peptides, only VQIVYK has a small net positive charge (Table 1) and thus likely binds stronger to the negatively charged citrate-coated NP surface, while GNNQQNY is uncharged and Aβ_40_ has an overall negative charge. The stronger the peptide-NP surface attraction, the stronger are the expected effects on peptide aggregation.

**Figure 2.**
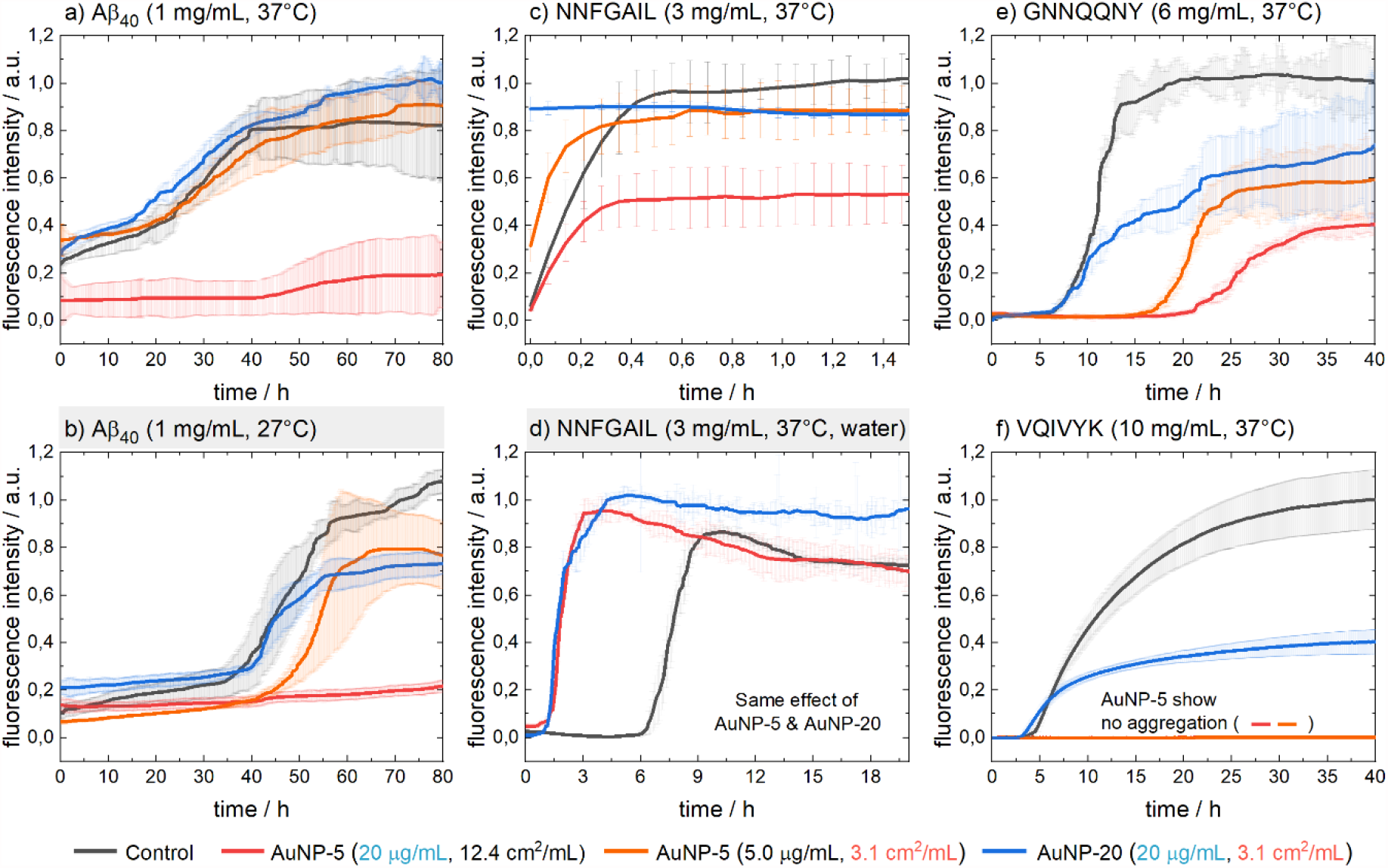
ThT fluorescence measurements to follow the aggregation kinetics of the peptides (a-b) Aβ_40_, (c-d) NNFGAIL, (e) GNNQQNY and (f) VQIVYK. Peptide aggregation was probed at 37°C in HEPES (100 mM, pH 7.4) buffer without and with gold NPs (AuNP-5, AuNP-20) present. In addition, highlighted in light-grey background, Aβ_40_ was studied at 27°C (b) and NNFGAIL in water (d) to probe the effects of temperature and solution conditions. An increase in ThT fluorescence indicates the aggregation of peptide monomers into fibrils. An overview of the half times of aggregation (t_1/2_, 50%) is summarized in the Supporting Information (Table S3).

As Aβ_40_ aggregation was not influenced by AuNP-5 at 5 µg/mL at 37°C, an additional experiment at 27°C was performed (Figure 2 b). At lower temperature, peptide aggregation was slower and NP surface effects became more relevant, as AuNP-5 now also retarded peptide aggregation at lower NP concentration. The larger NPs (AuNP-20) had slightly accelerating but no significant effects on the aggregation of the peptides Aβ_40_, GNNQQNY and VQIVYK (Figure 2 a, b, e, f). In contrast, NNFGAIL aggregated much faster than the other peptides (Figure 2 c), likely due to the high hydrophobicity and thus low solubility and high aggregation propensity (Figure S4). Both AuNP-5 and AuNP-20 accelerated NNFGAIL aggregation while AuNP-5 led to a lower yield of fibrils. As NNFGAIL immediately aggregated in the presence of AuNP-20 under the studied conditions (Figure 2 c), peptide aggregation was additionally studied in water (Figure 2 d). In unbuffered aqueous solution, both AuNP-5 and AuNP-20 strongly accelerated NNFGAIL aggregation; however, the effects of NPs could not be clearly distinguished due to pH changes caused by the addition of NP solutions that were citrate-stabilized (Table S2). Thus, it is essential to probe NP effects on peptide aggregation under buffered conditions, as done in this study. While Aβ_40_, GNNQQNY, and NNFGAIL in water (Figure 2 a, b, d, e) present typical sigmoidal nucleation-polymerization kinetics, VQIVYK and NNFGAIL in HEPES buffer (Figure 2 c, f) show an exponential fibril growth, indicating immediate aggregation seeded by already formed fibrils for NNFGAIL, and after a lag time for VQIVYK.^45^ The presence of AuNP-5 most likely inhibited the nucleation into oligomeric species and thus larger fibrils for Aβ_40_, GNNQQNY and VQIVYK (Figure 2 a, b, e, f). While both AuNP-5 and AuNP-20 provided a seeding surface for NNFGAIL, this was particularly apparent when studied in water and thus no electrostatic contributions from buffer salt (Figure 2 c, d).^38,46^

Dynamic light scattering (DLS) measurements were used to follow Aβ_40_ fibril growth *in situ* without the presence of a fluorescence dye at 37°C (Figure 3 a, c, e). The method works best for samples that only contain spherical particles of the same size that do not precipitate and follow perfect Brownian motion. Amyloid fibrils do not satisfy these assumptions; however, the measured hydrodynamic diameter grows when fibrils form and elongate. While the DLS hydrodynamic diameter of the peptide species increased in the 0-500 nm range in all cases, larger aggregates in the 2000-4000 nm range were only observed without AuNP and with AuNP-20 present, indicating an inhibition of the formation of mature peptide fibrils when AuNP-5 were present. In addition, a larger portion of mature fibrils was measured when AuNP-20 were present, compared to the sample without AuNPs. Despite the differences in time scale of aggregation between ThT fluorescence (Figure 2) and DLS (Figure 3) due to differences in the experimental setups (e.g. no shaking of DLS measurement cell), an inhibitory effect on Aβ_40_ aggregation by smaller NPs and an accelerating effect by larger NPs was confirmed.

**Figure 3.**
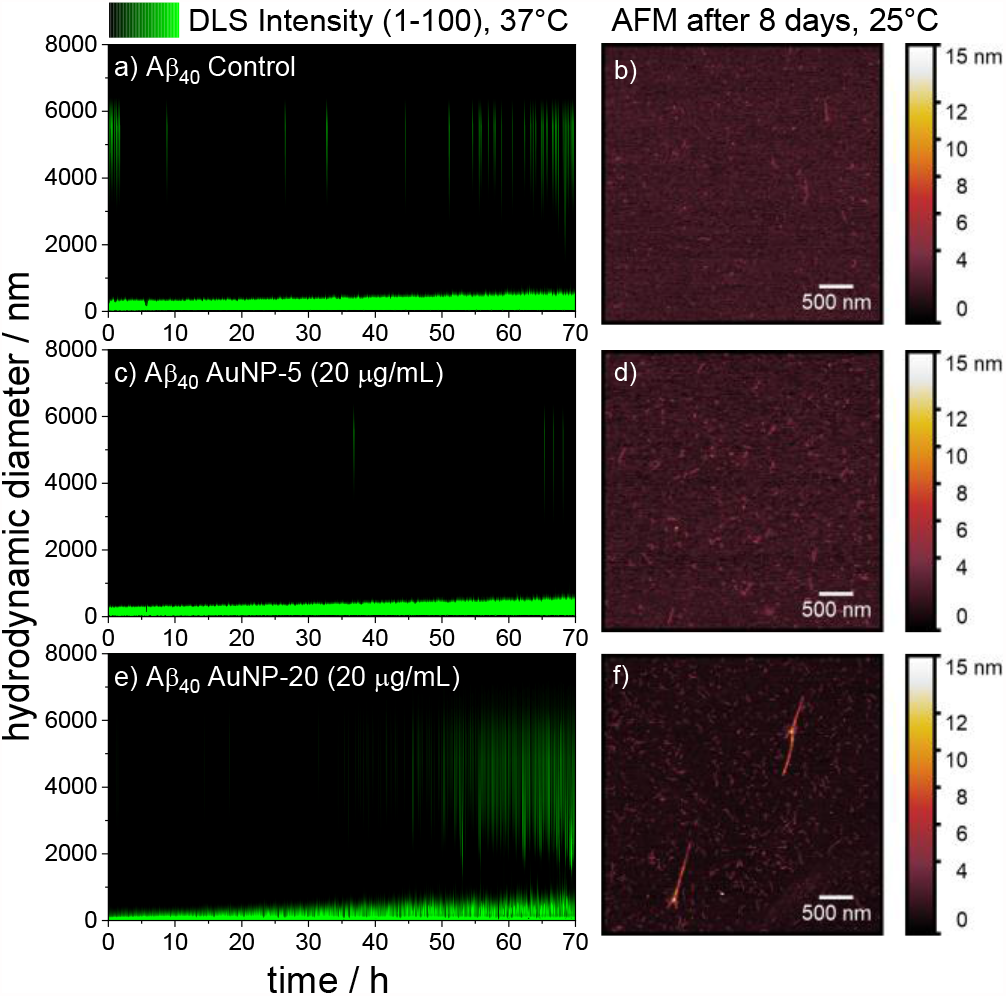
The kinetics of Aβ_40_ aggregation was followed using (a, c, e) dynamic light scattering (DLS) and independently (b, d, f) atomic force microscopy (AFM). Peptide aggregation was studied in HEPES buffer (100 mM, pH 7.4) without and with 20 µg/mL gold NPs (AuNP-5, AuNP-20) present. An increase in DLS hydrodynamic diameter indicates the formation of larger peptide aggregates. Note that small amounts of any particles in a pure water or buffer solution scatter light very strongly (Figure S6); thus, the analysis is focused on the formation of larger aggregates > 1000 nm in hydrodynamic diameter. AFM enabled the imaging of typical fibril structures after eight days. Additional AFM and scanning electron microscopy (SEM) images can be found in the Supporting Information (Table S4).

Atomic force microscopy (AFM, Figure 3 b, d, f) and scanning electron microscopy (SEM, Table S4) images of Aβ_40_ samples that were incubated for eight days at 25°C show that amyloid fibrils are formed both with and without NPs present. However, larger fibril structures were only observed in the samples without any NP and with AuNP-20 present (Figure 3 b, d, f and Table S4), while they were more prevalent in the sample with AuNP-20. Transmission electron microscopy (TEM) characterization of Aβ_40_, GNNQQNY and VQIVYK confirmed fibril formation in all cases following significant incubation times of up to two months (Figure 4, Table S5). While there were morphological differences between the peptides, TEM revealed more flexible amyloid fibril clusters in the presence of AuNP-5, indicating a structural influence of NP geometry on fibril morphology. The resulting Aβ_40_ fibrils varied between 11 and 14 nm in diameter, with finer and more flexible, yet densely packed, fibrillar structures in the presence of small diameter NPs. This trend was also observed for GNNQQNY in the presence of AuNP-5 (Figure 4). Thus, small NPs may additionally impact the peptide self-assembly as they are in the same size regime as the peptide fibrils.

**Figure 4.**
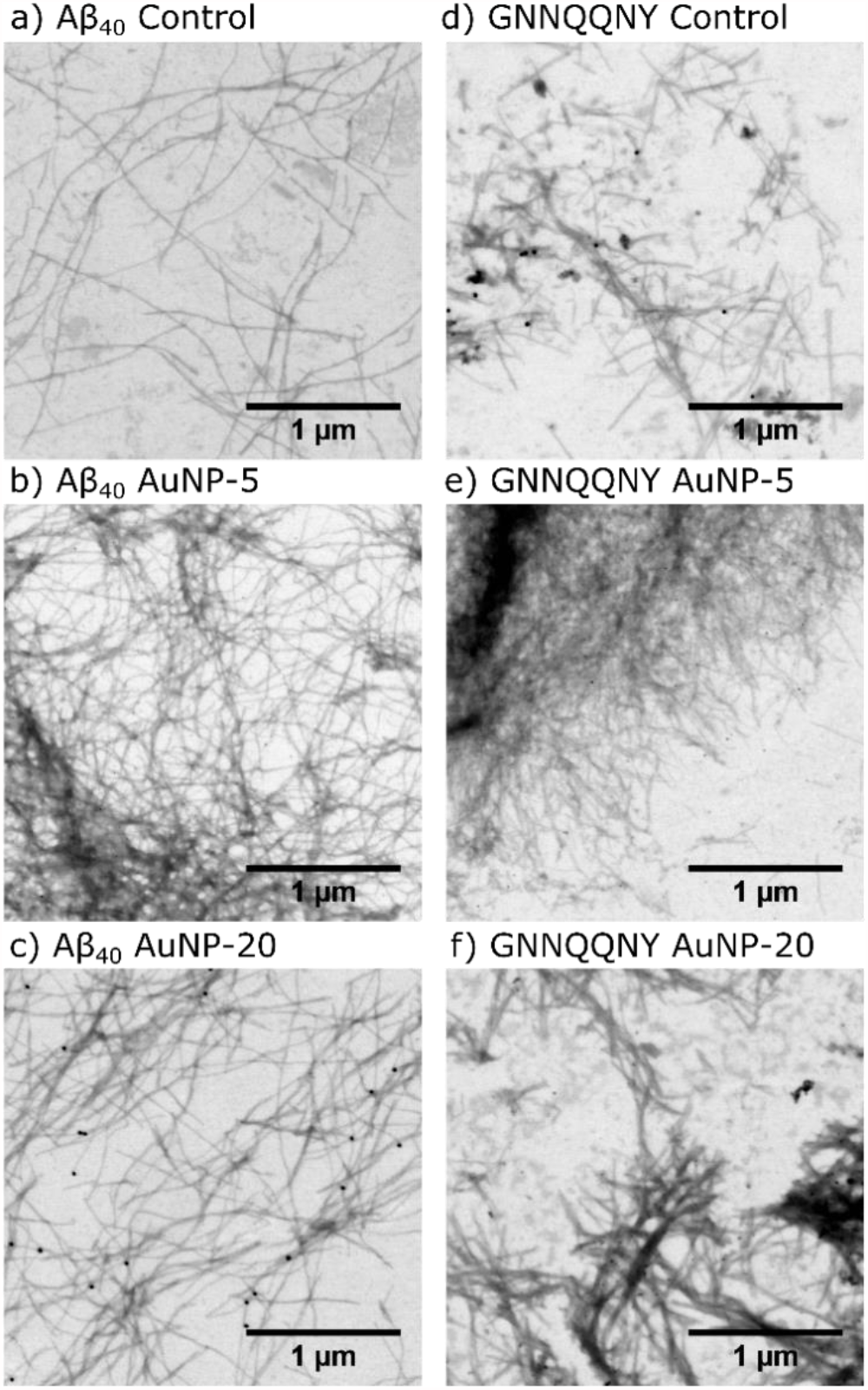
TEM images of (a-c) Aβ_40_ and (d-f) GNNQQNY peptide fibrils deposited onto silicon and air-dried after incubation in HEPES buffer without and with 20 µg/mL AuNP-5 and AuNP-20 for two months. Additional TEM images can be found in the Supporting Information (Table S5).

Molecular dynamics (MD) simulations were used to explain the influence of surfaces and curvature on amyloid fibril formation. We started with simulations of the interaction of the peptides at a planar citrate coated gold surface, using an atomistic gold model based on the GolP (Gold-Protein) force field, as previously reported (Figures 5 and S7).^5,9,47^ These simulations revealed that the conformational space sampled by the larger Aβ_40_ peptide was reduced if in contact with the surface, while the smaller NNFGAIL or GNNQQNY peptides were not restricted in flexibility (see Schlitter entropy in Figure 5 e). The shorter peptides remained bound via their N-terminus in most cases over the course of the simulations whereas the larger Aβ_40_, with either a β-sheet or an α-helix initial structure, desorbed or formed multiple interaction sites respectively, via the N-terminus as well as the lysine side chain at position 28 (K28). Once additional peptides were introduced into these systems, oligomeric structures were formed at the surface.^5^ Peptide-peptide interactions become more important, leading to more β-sheets (see Table S6 and Figure S8) as well as cluster formation and phase separation rather than surface binding (see Figures 5 f-g and S7 e-j). Models of the mechanistic influence of surfaces on peptide nucleation and aggregation have previously been reported and summarized.^5,9,30,48^

**Figure 5.**
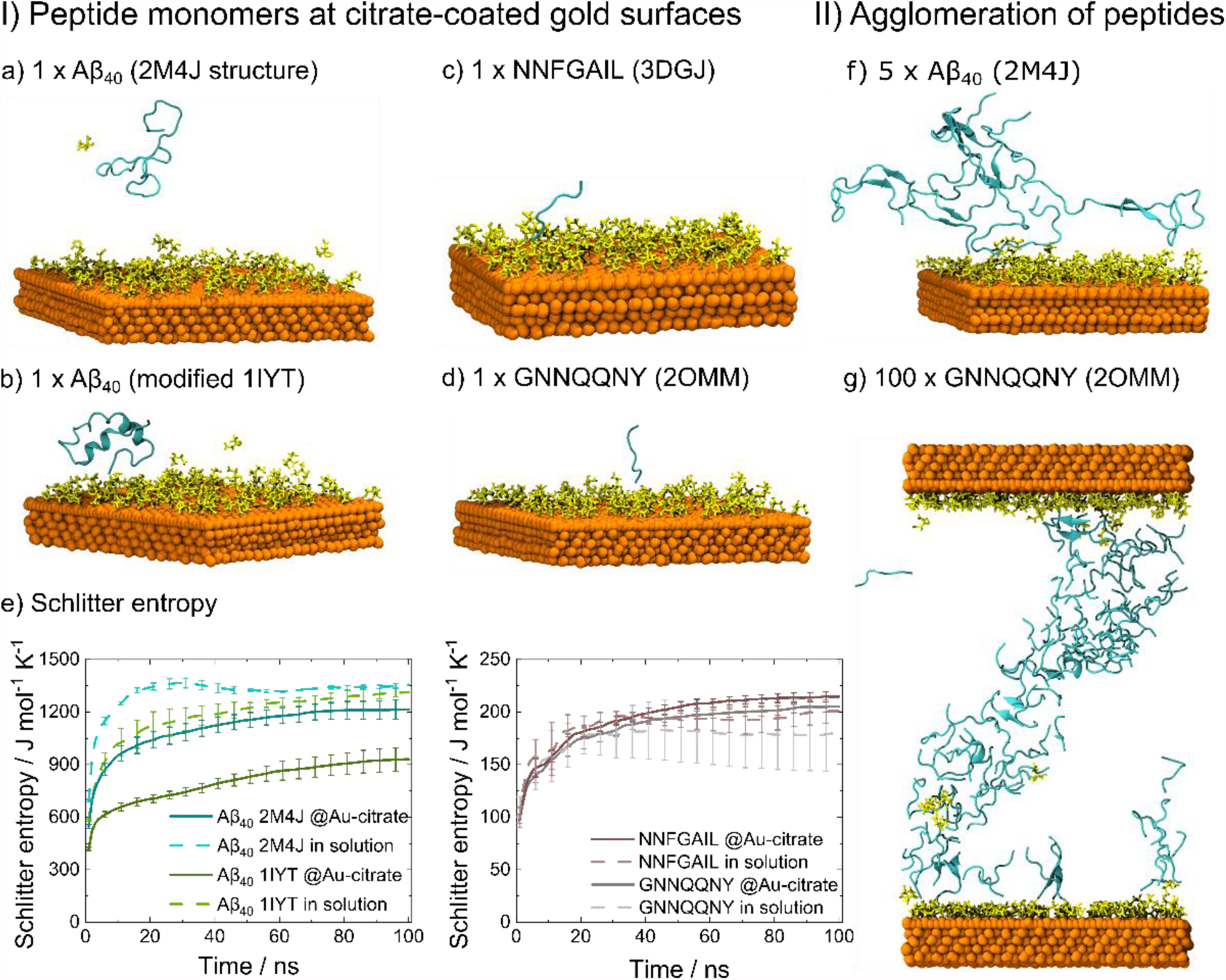
(I) MD simulation snapshots of peptide monomers of (a, b) Aβ_40_, (c) NNFGAIL and (d) GNNQQNY at citrate-coated (shown in yellow) gold surfaces. One bound peptide (a-d) was simulated for 100 ns in multiple repetitions to study conformational changes. (e) The conformational entropy was analyzed using the Schlitter method (C_α_ atoms). The larger Aβ_40_ peptide had multiple interaction sites with the surface when bound while the smaller NNFGAIL and GNNQQNY peptides were adsorbed via their N-terminus only, resulting in a reduction in conformational space for the Aβ_40_ peptide when bound with gold compared to in solution. Both (a) β-sheet (2M4J) and (b) α-helical (modified 1IYT) starting structures were studied for Aβ_40_. (II) When more peptide monomers were added and thus the peptide locally concentrated (f-g), peptide-peptide interactions became more important, inducing cluster formation, eventually resulting in percolated clusters that bridge the periodic simulation box (see in g).

To understand how intrinsic differences in surface geometry affect the stability of NP-bound peptide aggregates, additional MD simulations were performed. Peptide aggregates consisting of five or ten monomers aligned in a parallel β-sheet were studied, using fibril structure information from the PDB databank^41,49,50^ and representing the size range of critical nuclei for fibril growth.^51,52^ NP curvature was implicitly modelled using spherical harmonic potentials acting on the peptides’ N-termini (see Methods section for details). The effective stiffness of the interface of the NP was modeled by the choice of the force constant which we derived from the simulations of peptides at planar citrate-coated gold surfaces (see Figure 5). These suggested that the peptide N-termini favorably bind to the negatively charged citrates.^9,24^ Representations of the peptide oligomers of Aβ_40_, NNFGAIL and GNNQQNY after 100 ns are shown in Figure 6 (for 10mers; see Figures S9 and S10 for 5mers).

**Figure 6.**
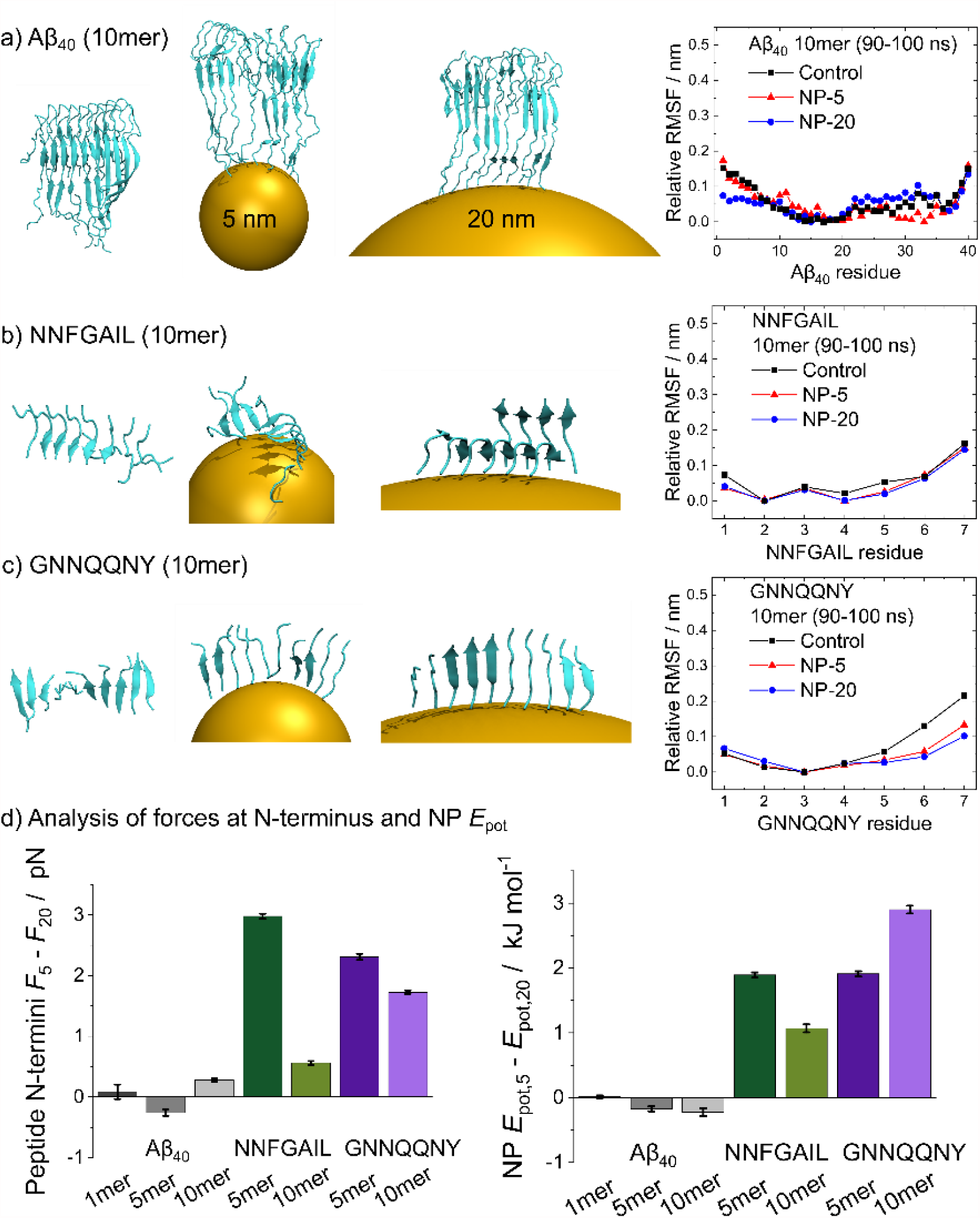
Oligomers of (a) Aβ_40_, (b) NNFGAIL and (c) GNNQQNY were studied when bound to implicit nanoparticles of 5 nm (NP-5) and 20 nm (NP-20) diameter and when in solution (Control). The relative RMSF (root-mean-square fluctuation, relative = minimum for each condition set to 0 to correct for the overall translational movement) of the peptides during the last 10 ns of simulation time is shown. Higher RMSF values indicate larger structural changes. (d) The NP curvature effect on peptide aggregates was analysed by quantifying the average absolute forces on the N-termini and the NP potential energy over the last 90 ns (10-100 ns) of simulation time. Relative differences between NP-5 and NP-20 are shown. The absolute values are largely curvature independent and determined by translational restraining and Brownian forces (see Table S8 for absolute values).

While the Aβ_40_ aggregates (Figure 6 a) show only subtle differences in peptide structure between the simulations in solution and those bound to implicit NPs, aggregates formed from the shorter heptapeptides NNFGAIL and GNNQQNY (Figure 6 b-c) rearranged on the NP surface. This is most likely due to the large, flexible structure of the Aβ_40_ peptide compared to the smaller heptapeptides. We emphasize that peptide aggregation occurs on time scales of several hours under experimental conditions, whereas simulations can only examine the initial changes in structure. The β-sheet content of the peptides increased in most simulations compared to the starting structure (see Table S7). NNFGAIL formed β-sheet double layers, indicating the stability of fibril oligomers when bound to curved surfaces, especially with NP-20. Further analysis of the relative RMSF (root-mean-square fluctuation) of the peptide aggregates confirmed a high structural stability during the last 10 ns of simulation time, while larger structural rearrangements particularly within the N-terminal region of Aβ_40_ were observed during the initial steps (Figure S9).

The curvature stress imposed on the aggregates has been investigated further by quantifying the average absolute forces that act on the peptides’ N-termini and the resulting potential energies to restrain the peptides on the implicit curved NP surfaces (Figure 6 d). Since both measures are largely curvature independent and result from restraining of translational motion and Brownian forces, the relative difference between the effects at NP-5 and NP-20 were calculated (see also Table S8 for absolute values). While the difference between NP-5 and NP-20 was not significant for Aβ_40_, possibly due to Aβ_40_’s large size, the forces on the N-termini of NNFGAIL and GNNQQNY aggregates were 0.5-3 pN higher for the NP-5 in all cases. Thus, peptide fibril aggregates were energetically less favored at higher curved surfaces. The observed stress imposed by the curvature can explain why similar CFGAILSS peptide motifs showed decreased β-sheet formation at higher surface curvatures.^27^ Finally, since stress imposed by the curvature drives structural destabilization of the aggregates, and given its observed magnitude of only a few pN on each peptide, structural adaptions such as loss of β-sheet content would require larger timescales of microseconds or more which were computationally not feasible. Owing to the greater degree of flexibility within the Aβ_40_ tail, surface curvature destabilization was not sampled in our model simulations. Differential curvature effects could affect stabilization of Aβ_40_ oligomers larger than the 10mer used in our study.

It is important to emphasize that NP curvature, even when extreme, does not significantly affect the adhesion strength and residence time of peptides on the surface of NPs. For example, in the case of NFFGAIL, as a result of thermal fluctuations, the binding force observed in the equilibrium simulations on planar citrate-coated gold withstands values of up to 48 *k*_B_ *T* nm^-1^ or about 200 pN. In comparison, the excess force due to curvature coupling, i.e. the difference between 5 nm and 20 nm NPs, is two orders of magnitude smaller and is only in the low pN range. Hence, the absolute surface force measured in our implicit NP simulations, being about 20 pN, is predominantly due to spatially restraining the N-terminus and originates from Brownian forces (collisions with solvent) and well as the reduction of translational entropy. Simulations of Aβ_40_ monomers at curved surfaces (implicit NPs with diameters of 5 nm, 20 nm, and 1 mm – as a planar model) revealed that curvature did also not significantly impact the configurational entropy in our model simulations (see Figure S11). Since reversible (un)binding events are rare on the timescale of our simulations, and since surface curvature is not expected to significantly affect (un)binding frequencies, we effectively modelled the peptide N-terminus as covalently bound.

Our simulations demonstrated that surface curvature is capable of inducing stress towards β-sheet rich peptide oligomers which in turn inhibits amyloid fibril formation. This NP inducible stress depends on the degree of NP curvature as well as the intrinsic aggregation propensity and binding affinity of the peptide to NP surfaces. The same material can result in a variety of interactions with biomolecules, as previously observed for convex and concave curved surfaces of carbon nanotubes.^53^ Arkin and Janke theoretically studied a wide range of curvatures for non-grafted and end-grafted polymers on NPs.^28^ Our implicit NP curvature simulations modelled a scenario where all peptide N-termini remained bound to the NP surface. However, in nature, the impact of NPs also results from a competition between peptide-peptide and peptide-NP attraction, with the latter being dependent on the NP surface and the specific properties of each peptide.^5^ The surface binding affinity of a peptide is thus an additional contribution to the NP effect, and differences in amino acid sequence and their affinity towards binding a NP of varying size might lead to different structural motifs and outcomes.^54^

In this work, we have provided an initial model that explains the complex effects of surface curvature on amyloid fibril formation. While larger NPs provided an attractive surface for seed formation and oligomer growth, smaller NPs interfered with the structure of fibrils and destabilized protofibril oligomers; thus inhibiting amyloid peptide aggregation. Moreover, peptide-NP attraction needed to be in an intermediate range that enables conformational changes, influenced by the peptide sequence and solution ionic strength, as has previously been reported.^5,9^ By studying multiple peptides, we illustrated that the NP size effect is not the same or universal for each peptide but depends on the peptide-surface attraction. Too strong surface attraction can remove significant amounts of peptide from solution, leading to amorphous aggregates, thereby preventing fibril formation. To compensate the adsorption potential resulting from such nanostructures, surfaces may be tailored with coatings, as has been extensively reported.^21,32,55,56^ This is particularly relevant for surfaces with designed function for applications in biomedicine and nanotechnology.^57^

## CONCLUSIONS

The curvature effects of nanostructures on peptide oligomer structure and subsequently amyloid peptide aggregation presented in this study were rationalized based on experimental observations and theoretical models. High NP curvature caused stress in fibril oligomer structures, destabilized those and potentially caused inhibitory effects on fibril formation as well as changes in fibril morphology, while low NP curvature provided a seed surface. Our findings may provide an approach for the study of biomolecule conformations that are restrained on curved surfaces. In nature, materials are often not ideally spherical or planar but typically small defects can lead to a number of additional nanoscale substructures that will govern the interaction with biomolecules. The complex self-assembly behavior of amyloidogenic peptides thus needs to be always considered in the context of its surrounding physicochemical environment and present interfaces.

## METHODS

### Peptides and Buffer Preparation

The peptides amyloid beta (Aβ_40_, purity 98%), GNNQQNY (99%), VQIVYK (99%) and NNFGAIL (98%) were synthesized and purified by the Core Unit of Peptide Technologies at Leipzig University (Leipzig, Germany). In addition, Aβ_40_ (95%) was obtained from Peptide 2.0 (Chantilly, VA), and NNFGAIL (96%) from Eurogentec (Seraing, Belgium). Dimethyl sulfoxide (DMSO, ≥99.9%) was obtained from Merck (Darmstadt, Germany). Sodium phosphate monobasic (≥99.0%) was obtained from Sigma-Aldrich (St. Louis, MO). HEPES (≥99.5%) and TRIS (≥99.9%) were purchased from Carl Roth (Karlsruhe, Germany). Ultrapure water (18.2 MΩ cm) was used for all experiments (Sartorius, Göttingen, Germany, and Merck, Darmstadt, Germany). Sodium hydroxide and hydrochloric acid were used to adjust the pH to 7.40±0.05. HEPES buffer was used at a final concentration of 100 mM.

### Gold Nanoparticle Synthesis

#### AuNP-5, AuNP-20

AuNPs (citrate coated) in sizes 5 nm (AuNP-5, EM.GC5, batches 19070110, 019965 and 026755) and 20 nm (AuNP-20, EM.GC20, batches 20060186, 021195 and 026158) were purchased from BBI Solutions (Cardiff, UK). AuNPs were delivered as suspensions in water with no preservative residual chemical left from manufacture. The solutions had an orange-red (AuNP-5, 63.2 µg/mL) and red (AuNP-20, 56.6 µg/mL) color.

#### AuNP-mix

AuNPs (citrate coated) were synthesized using a chemical reduction method (Frens *et al*.)^58^ by heating up 10 µL HAuCl4 (99.99% trace metals basis, 30 wt. % in dilute HCl, Sigma-Aldrich, St. Louis, MO) in 29.99 mL ultrapure water (0.5 mM) to its boiling point. 840 µL trisodium citrate (34 mM, 1 wt. %, ≥99%, ABCR, Karlsruhe, Germany) were quickly added to the solution to yield a AuNP solution, as reported previously.^59^ The solution was stirred for 30 minutes. A dark red solution of AuNP-mix (92 µg/mL) was obtained.

Experimental details of the characterization of the AuNPs by dynamic light scattering (DLS), zeta potential measurements and UV-vis absorption spectroscopy are provided in the Supporting Information (Table S1 and Figure S1).

### Thioflavin T Fluorescence Assays

Fibrillation kinetics were measured using fluorescence spectroscopy. Thioflavin T (ThT, Sigma-Aldrich, St. Louis, MO) was used as a fluorescent dye which shows increased fluorescence intensity at 482 nm when bound to fibrils showing the cross-β-structure.^44^ ThT was diluted in DMSO to obtain a stock solution that was stored at −20°C and protected from light. The final concentration of ThT in the assay was 20 μM. All peptides were first dissolved in pure DMSO (50 µL per 1 mL final volume) before the stock solution was diluted in HEPES buffer (100 mM, pH 7.4) or water. The nanoparticle solutions were added as a final step. The total sample volume in each well was 150 µL. Each experiment was repeated at least in triplicate. Note that the described order of the addition of components to the sample is important. When the peptide stock solution was added last, peptide aggregation started immediately due to a temporarily high peptide concentration exposed to both buffer and nanoparticles, which should be avoided.

Black polystyrene 96 well microplates with solid, flat bottom (Nunc, Thermo Fisher Scientific, Waltham, MA) were used. All empty wells were filled with water to avoid drying of the most outer wells over time. The ThT fluorescence was recorded using a Tecan infinite M200 microplate reader (Tecan, Männedorf, Switzerland) with excitation and emission wavelength set to 440/9 nm and 482/20 nm, respectively. The fluorescence intensity was measured every five minutes with two seconds of shaking (2 mm linear shaking amplitude) prior and after each measurement. The experiments were performed at 27 and 37°C.

The NNFGAIL experiment in water (Figure 2d) was performed in black polystyrene 96 well microplates with solid, clear bottom and non-binding coating (Greiner Bio-One, Kremsmünster, Austria) with ThinSeal™ adhesive sealing films (Astral Scientific, Taren Point, Australia). The ThT fluorescence was recorded using a CLARIOstar plate reader (BMG Labtech, Ortenberg, Germany) with excitation and emission wavelength set to 440/10 nm and 480/10 nm, respectively. The microplate was agitated for 40 seconds before each measurement cycle (five minutes) using double orbital shaking (300 rpm). The experiment was performed at 37°C. Note that experiments with NNFGAIL in HEPES buffer using this set-up led to consistent accelerating effects of the NPs on NNFGAIL aggregation.

Averaged data of the repetitions with standard error bars were smoothed using parameters that removed signal noise and plotted with OriginPro 2021 (OriginLab Corp., Northampton, MA). The highest fluorescence intensity measured for each peptide was normalized to zero. For the NNFGAIL experiment in water (Figure 2d), initial fluorescence values were set to 0, to be consistent with the instrument sensitivity of all other experiments. Half times of aggregation were determined as times at which the half-maximum fluorescence (*t*_*1/2*_, 50%) was reached, in respect to the baseline fluorescence intensity.

### Dynamic Light Scattering (DLS) to Follow Peptide Aggregation

DLS is based on the autocorrelation of a measured intensity of time curve of backscattered laser radiation. Larger particles move slower and thus, the exponential decay of the autocorrelation is slower as well. The particle size is reconstructed from these information.^60^ Measurements were performed on a Zetasizer Nano ZSP instrument (Malvern Instruments, Malvern, UK) in polystyrene disposable cuvettes at 37°C. Aβ_40_ peptide (1 mg/mL, 50 µL DMSO per 1 mL solution) without and with AuNP-5 or AuNP-20 present (20 µg/mL) was studied in HEPES buffer (0.1 M, pH 7.4). Samples were continuously incubated without shaking in the measurement cell over time. DLS was recorded at a wavelength of *λ* = 633 nm and at a scattering angle of 173° for a duration of 180 s. Dispersant was water (viscosity 0.8872 mPas, refractive index 1.330, and dielectric constant 78.5) and sample material was protein (refractive index 1.45, absorption 0.001). Data were plotted with OriginPro 2021 (OriginLab Corp., Northampton, MA).

### Atomic Force Microscopy (AFM)

Silicon wafers (G3390, Plano, Wetzlar, Germany) were used as substrates to measure peptide fibrils. The wafers were cleaned in ethanol (99.8%, Carl Roth, Karlsruhe, Germany), gently shaken and dried under a stream of nitrogen gas prior to use. Aβ_40_ samples were deposited onto the substrates by dropping 5 µL sample solution on the silicon wafers for five minutes before the wafers were gently rinsed with 1 mL ultrapure water and dried under a stream of nitrogen gas. Aβ_40_ samples (1 mg/mL, 50 µL DMSO per 1 mL solution) without and with AuNP-5 or AuNP-20 present (20 µg/mL) were incubated at 25°C in HEPES buffer (0.1 M, pH 7.4). Samples were shaken every 5 min for 4 s (200 rpm), similar to the ThT fluorescence assay, and 1:9 v/v diluted samples (100 µL sample, 900 µL HEPES buffer) were measured at multiple time points. AFM measurements were performed on a Dimension Icon instrument (Bruker, Billerica, MA) with an OTESPA R3 tip (Bruker, Billerica, MA) in tapping mode at room temperature in air. The nominal spring constant of the cantilevers was 26 N/m and the nominal frequency 300 kHz. AFM images were processed using Nanoscope 8.15r3 and Gwyddion 2.51 (http://gwyddion.net/).^61^ For processing, rows on the images were aligned, the base flattened and the background levelled using a polynomial fit, if needed.

### Transmission electron microscopy (TEM)

The morphologies of Aβ_40_, GNNQQNY and VQIVYK fibrils were verified using TEM. Sample solutions from ThT fluorescence measurements (different concentration to the ones presented in the manuscript) were diluted 1:38 with pure water after two months (incubated in a fridge to enable a slow formation and/or growth of fibrils after ThT measurement). 1 µL droplets of these solutions were applied on formvar coated copper grids, allowed to dry for at least 1 hour and negatively stained with 1% uranyl acetate in pure water. Transmission electron micrographs were recorded using a Zeiss SIGMA electron microscope (Zeiss NTS, Oberkochen, Germany) equipped with a STEM detector and Atlas Software. TEM images were processed using ImageJ 1.52a (https://imagej.nih.gov/ij/).^62^

### MD Simulations of Planar Citrate-Coated Gold Surfaces

The interaction of amyloidogenic peptides with citrate-coated gold surfaces was simulated using GROMACS 4.5.7.^63–67^ The MD simulations were performed following a method previously published.^5^ Input structures for NNFGAIL (PDB: 3DGJ),^41^ GNNQQNY (PDB: 2OMM),^50^ Aβ_40_ (β-sheet, PDB: 2M4J)^49^ and Aβ_42_ (β-sheet, PDB: 2NAO)^68^ were selected from the Protein Data Bank (PDB). An α-helical structure of Aβ_42_ (PDB: 1IYT)^69^ was adapted for Aβ_40_ to simulate an alternative secondary starting structure. The peptides GNNQQNY and NNFGAIL became unstructured as monomers. N- and C-termini of the peptides were charged.

A planar, ten atoms thick, in xy continuous, gold surface was used to approximate the spherical nanoparticles. This approximation is particularly appropriate to study the effect of larger AuNPs (e.g. AuNP-20), as the curvature seen by a pair of peptides is very low.^9^ The Au(111) parameters are based on the GolP (Gold-Protein) force field^47^ and were used in an adapted form to correct for the citrate adsorption on the gold surface as previously described.^5,9^ Force field parameters for the adsorbed citrate anions were used from Brancolini *et al*.^70,71^

Periodic boundary conditions were applied. Peptides, water and ions were described with the OPLS/AA force field.^72,73^ Simulation parameters were used as previously published.^5^ The Particle Mesh Ewald method (PME) with a grid of 0.12 nm, a fourth order spline interpolation and a Coulomb cut-off at 1.1 nm was used to describe electrostatic interactions^74,75^ and a Lennard-Jones cut-off distance of 1 nm with a smooth switch off at 0.9 nm was used to describe van der Waals interactions. Interactions were updated every fifth step with a time step of 2 fs. Centre of mass motion was removed for the system (except gold) at every step. All bonds were constrained to their equilibrium values using the LINCS algorithm.^76^ Explicit water (Simple Point Charge, SPC)^77^ was constrained using the SETTLE algorithm.^78^

All systems were built in an orthorhombic simulation box (8.12 × 6.446 × 15 nm^3^) with the gold frozen during the simulations. The cell size was chosen so that periodic image interactions from neighboring boxes are prevented, unless this was intended (Figure 5 f-g). The systems were solvated and 150 mM sodium chloride was added as a physiological salt and to electro-neutralize the solutions. A steep concentration gradient was used to add positively charged sodium ions with a maximum close to the negatively charged citrate-coated gold surface. This was achieved using a slab geometry with Ewald summation correction to prevent electric fields through the box.^9,79^ After energy minimization using a steepest-deepest algorithm, the systems were run in a NVT ensemble. The velocity rescale algorithm was applied to couple the temperature to 300 K.^80^

Simulations of the peptides GNNQQNY and NNFGAIL with up to 30 randomly placed monomers each were previously reported.^5,9^ Starting from these simulations, the peptide GNNQQNY was further investigated at very high concentrations to mimic a local concentration of the peptide. Peptide monomers were added in steps (30+5 for 10 ns, 35+5 for 20 ns, 40+5 for 20 ns, 45+5 for 20 ns, 50+5 for 20 ns, 55+5 for 20 ns, 60+5 for 20 ns, 65+5 for 20 ns, 70+30 for 150 ns in triplicate) until reaching 100 peptide molecules (details in Table 2). The peptides Aβ_40_ and Aβ_42_, each with β-sheet and α-helix starting structures, were simulated with five randomly placed peptide monomers outside the gold-citrate layer (each once for 200 ns). From the above simulations, three surface-bound peptide monomers were chosen for the peptides GNNQQNY, NNFGAIL, and Aβ_40_ (α-helix and β-sheet). The surface bound peptides were simulated three times with different starting velocities (repetitions) for 100 ns to study their conformational flexibility, i.e. each peptide (GNNQQNY, NNFGAIL, Aβ_40_ α-helix, Aβ_40_ β-sheet) was simulated nine times (three different starting configurations, each with three different starting velocities). In addition, each system was run in water without gold surface for 100 ns in triplicate. All simulations are summarized in Table 2.

**Table 2.** Overview of MD Simulations with Planar Citrate-Coated Gold Surfaces.

| Peptide and sequence | Starting configuration and structure | Peptide concentration<sup>[a,b]</sup> | Simulation length |
| --- | --- | --- | --- |
| Aβ<sub>40</sub> | DAEFRHDSGYEVHHQKLVFFAEDVGSNKGAIIGLMVGGVV | | |
| | 5 peptides @ Au / β-sheet 2M4J<sup>49</sup> | 12 mM / 53 mg/mL | 1 x 200 ns |
| | 5 peptides @ Au / α-helix 1IYT<sup>69</sup> | 12 mM / 53 mg/mL | 1 x 200 ns |
| | 1 peptide @ Au / β-sheet 2M4J<sup>49</sup> | 2.4 mM / 10.5 mg/mL | 3 x 3 x 100 ns |
| | 1 peptide in solution / β-sheet 2M4J<sup>49</sup> | 2.8 mM / 12 mg/mL | 3 x 100 ns |
| | 1 peptide @ Au / α-helix 1IYT<sup>69</sup> | 2.4 mM / 10.5 mg/mL | 3 x 3 x 100 ns |
| | 1 peptide in solution / α-helix 1IYT<sup>69</sup> | 2.8 mM / 12 mg/mL | 3 x 100 ns |
| Aβ<sub>42</sub> | DAEFRHDSGYEVHHQKLVFFAEDVGSNKGAIIGLMVGGVVIA | | |
| | 5 peptides @ Au / β-sheet 2NAO<sup>68</sup> | 12 mM / 55 mg/mL | 1 x 200 ns |
| | 5 peptides @ Au / α-helix 1IYT<sup>69</sup> | 12 mM / 55 mg/mL | 1 x 200 ns |
| NNFGAIL | NNFGAIL |  |  |
|  | 1 peptide @ Au / unstructured<sup>[c]</sup> | 2.4 mM / 1.8 mg/mL | 3 x 3 x 100 ns |
|  | 1 peptide in solution / unstructured | 2.7 mM / 2.0 mg/mL | 3 x 100 ns |
| GNNQQNY | GNNQQNY |  |  |
|  | 30+5 peptides @ Au / unstructured<sup>[c]</sup> | 85 mM / 71 mg/mL | 1 x 10 ns |
|  | 35+5 peptides @ Au / unstructured | 97 mM / 81 mg/mL | 1 x 20 ns |
|  | 40+5 peptides @ Au / unstructured | 109 mM / 91 mg/mL | 1 x 20 ns |
|  | 45+5 peptides @ Au / unstructured | 121 mM / 102 mg/mL | 1 x 20 ns |
|  | 50+5 peptides @ Au / unstructured | 134 mM / 112 mg/mL | 1 x 20 ns |
|  | 55+5 peptides @ Au / unstructured | 146 mM / 122 mg/mL | 1 x 20 ns |
|  | 60+5 peptides @ Au / unstructured | 158 mM / 132 mg/mL | 1 x 20 ns |
|  | 65+5 peptides @ Au / unstructured | 170 mM / 142 mg/mL | 1 x 20 ns |
|  | 70+30 peptides @ Au / unstructured | 243 mM / 203 mg/mL | 3 x 250 ns |
|  | 1 peptide @ Au / unstructured | 2.4 mM / 2.0 mg/mL | 3 x 3 x 100 ns |
|  | 1 peptide in solution / unstructured | 2.7 mM / 2.3 mg/mL | 3 x 100 ns |

[a] Peptide concentrations were calculated based on the total volume of water molecules in the simulation box, when no or only one peptide molecule was present.
[b] Some peptide concentrations are higher than in the experimental studies. These simulations were performed to investigate how the peptides behave when locally concentrated. Further, these simulations were used to extract the configuration of bound monomers for subsequent simulations. The box size was kept constant for all simulations with gold.
[c] The structures for GNNQQNY (PDB: 2OMM)<sup>50</sup> and NNFGAIL (PDB: 3DGJ)<sup>41</sup> were obtained from the Protein Data Bank. However, as monomers, the peptides were used in unstructured form.

The conformational entropy of the peptides (C_α_ atoms) at the gold surface (@Au) and in solution was determined based on the Schlitter formula using the GROMACS tools g_covar and g_anaeig.^81^ Minimum distances between the gold surface and the amino acid C_α_ atoms of the peptides were determined for the last 10 ns of simulation time using the GROMACS tool pairdist, revealing which Aβ_40_ residues were bound most closely to the surface. The secondary structure content of the peptides was analyzed using the DSSP tool (Define Secondary Structure of Proteins, do_dssp) for the last 10 ns of simulation time.^82,83^ Averaged data of the repetitions are presented.

### Molecular Dynamics Simulations (MD) with Implicit NPs

#### a) Model

The curvature effect inherently imposed by NPs was modelled via external potentials that were implemented in the Gromacs 4.5.7 software.^63–67^ The implicit NP was constructed such that its surface exactly crosses the center of the simulation box (Figure 7). The NP itself was not subject to periodic boundary conditions (PBC). It should be emphasized that the AuNPs in the experiments were citrate capped and that physisorption of peptides mainly took place via N-terminal binding to the negatively charged citrates adhered on the NP rather than the gold surface itself.^5,9^ The relatively soft nature of such a physisorption is modelled by applying a spherical harmonic potential, *E(r-r*_*0*_) = 1/2·*k*·(*r-r*_*0*_)^2^, with *k* being a force constant (500 kJ nm^-2^ mol^-1^), *r*_*0*_ the effective radius of the NP, and *r* the position of the N terminal heavy atom with respect to the NP center. The remainder of the protein experiences the excluded volume imposed by the NP via a flat bottom potential, *E(r-r*_*0*_) = 1/2·*k*·(*r-r*_*0*_)^2^ if *r < r*_*0*_ and *E(r)* = 0 otherwise. Finally, the implicit NP was kept transparent for water and (counter) ions to mimic the hydrophilic nature of the particle’s surface (citrates, surface charge, ions and water). The NP radius *r*_*0*_ was simulated at 2.5 nm and 10 nm to study the respective NP-5 (diameter 5 nm) and NP-20 (diameter 20 nm).

**Figure 7.**
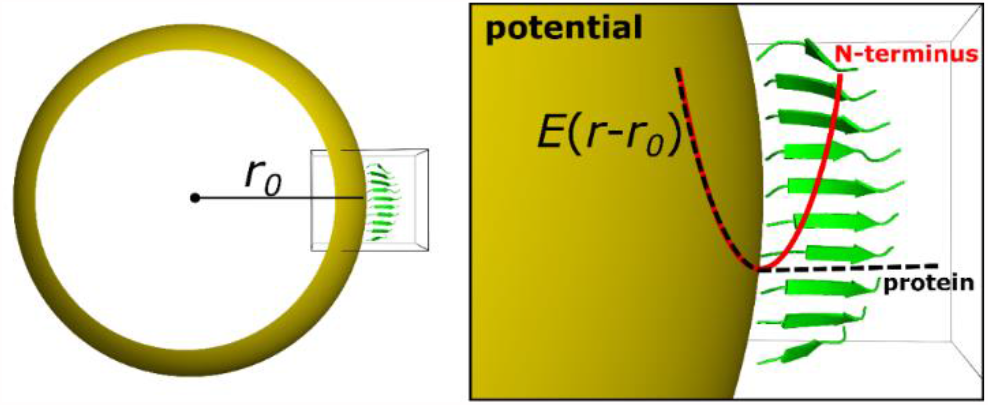
Schematic illustration of implicit NP simulations. NP (yellow) curvature was varied by changing the radius *r*_*0*_. A peptide β-sheet (green) is adsorbed to the NP surface with its N-terminus. A spherical harmonic potential *E*(*r*-*r*_*0*_) was used to model the interactions between peptide N-terminus and NP surface. The red line represents the adsorption (attractive potential) of the peptide N-terminus to the surface, whereas the black line mimics the repulsion of the protein when reaching into the excluded volume (*r* < *r*_*0*_).

The value of the force constant *k* of the NPs was estimated from the observed height fluctuations Δ*h* of the peptide N-terminus in the simulations of planar citrate-coated gold surfaces. To this aim, we constructed the probability distribution *P*(Δ*h*) from the trajectory of the simulations of the NFFGAIL peptide. The concomitant free energy profile is the Boltzmann inverse of *P*(Δ*h*), which is given by *F*(Δ*h*) = −*k*_*B*_·*T*·ln *P*(Δ*h*). Since the form of *P*(Δ*h*) is close to a Gaussian distribution, its Boltzmann inverse is an harmonic function of the form *F*(Δ*h*) = *k*·*x*^2^ + *c*_*1*_·*x* + *c*_*2*_, with *k* being the force constant of interest (Figure 8). We identified *k* to be roughly 1200 k_B_ T nm^-2^ or about 3000 kJ nm^-2^ mol^-1^. However, this value most likely provided an upper-bound since we cannot access the detailed nature of the electric double layer of NPs within experiments, i.e. additional ions or citrate layers, as well as the surface charge of gold NPs due to free electrons. Since the presence of additional electrostatic screening is expected to substantially ‘soften’ the apparent surface, we chose a more conserve value of 500 kJ nm^-2^ mol^-1^ to better describe the anticipated surface of NPs in experiments. It is important to emphasize that the excess force due to curvature coupling, i.e. the difference between 5 nm and 20 nm NPs, is only in the low pN range and is thus two orders of magnitude smaller than the instantaneous forces acting on the N-terminus due to the presence of thermal fluctuations (about 200 pN at Δ*h* = 0.05 nm in Figure 8). The forces acting on the absorbed peptides due to curvature coupling are thus very subtle and unlikely to lead to significant unbinding on the time scales of molecular simulations. Consequently, peptides can be modeled as covalently bound within our simulations.

**Figure 8.**
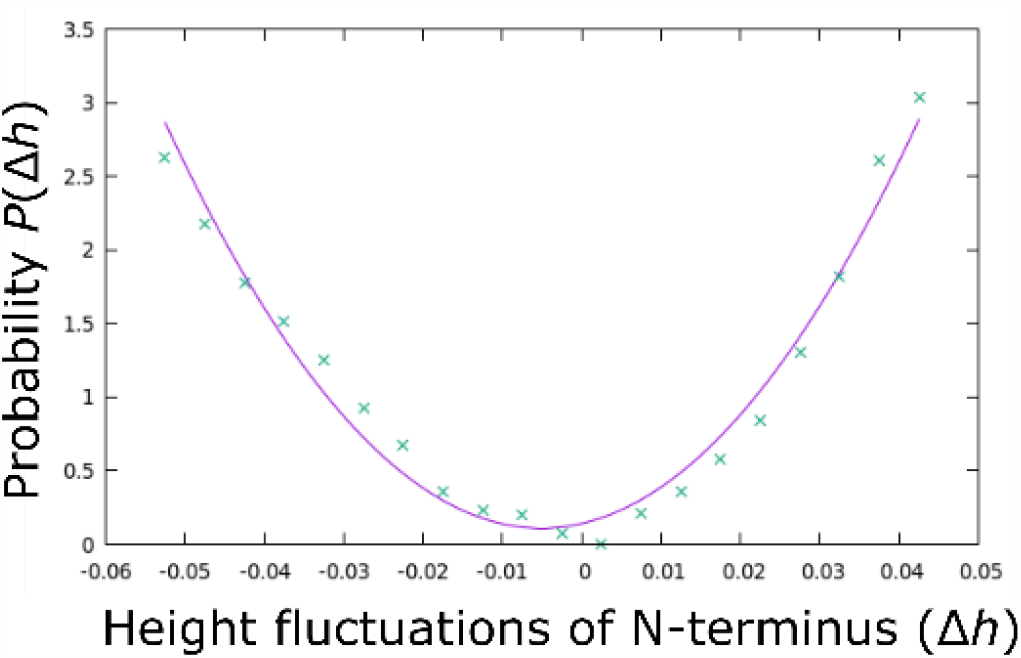
Probability distribution *P*(Δ*h*) of the observed height fluctuations Δ*h* of the peptide N-terminus in the simulations of planar citrate-coated gold surfaces of the NFFGAIL peptide.

#### b) MD Simulations

Simulation parameters were chosen as described above for the citrate-coated gold surfaces and are repeated here for completeness: Periodic boundary conditions were applied. Peptides, water and ions were described with the OPLS/AA force field.^72,73^ Simulation parameters were used as previously published.^5^ The Particle Mesh Ewald method (PME) with a grid of 0.12 nm, a fourth order spline interpolation and a Coulomb cut-off at 1.1 nm was used to describe electrostatic interactions^74,75^ and a Lennard-Jones cut-off distance of 1 nm with a smooth switch off at 0.9 nm was used to describe van der Waals interactions. Interactions were updated every fifth step with a time step of 2 fs. Centre of mass motion was removed at every step independently for “peptides” and “water & ions” in simulations with NP curvature present, or for the whole “system” in simulations without NP curvature present (and in 3 repetitions of Aβ_40_ simulations with NP curvature present). All bonds were constrained to their equilibrium values using the LINCS algorithm.^76^ Explicit water (Simple Point Charge, SPC)^77^ was constrained using the SETTLE algorithm.^78^ The systems were solvated and 150 mM sodium chloride was added as a physiological salt and to electro-neutralize the solutions. After energy minimization using a steepest-deepest algorithm, the systems were run in a NVT ensemble. The velocity rescale algorithm was applied to couple the temperature to 300 K.^80^

The stability of preformed β-sheets of the peptides Aβ_40_, NNFGAIL, and GNNQQNY (5mer and 10mer) were simulated at NP-5 (5 nm diameter) and NP-20 (20 nm diameter) surfaces. The peptide β-sheets were positioned close to the NP surface (central nitrogen atom of each oligomer was placed on implicit curvature surface). The simulation box sizes were chosen large enough so that the peptides did not cross the periodic box in the dimension of the curvature potential, with a minimum distance of 2 nm between peptide solute and box. Each simulation was run for 100 ns in triplicate (see Table 3 for details). Input structures for Aβ_40_ (β-sheet, PDB: 2M4J)^49^, NNFGAIL (PDB: 3DGJ)^41^ and GNNQQNY (PDB: 2OMM)^50^ were selected from the Protein Data Bank (PDB). N- and C-termini of the peptides were charged. VMD was used to visualize the simulations and GROMACS tools were used for analysis.^84^ Data were plotted with OriginPro 2017 and 2019 (OriginLab Corp., Northampton, MA).

**Table 3.**
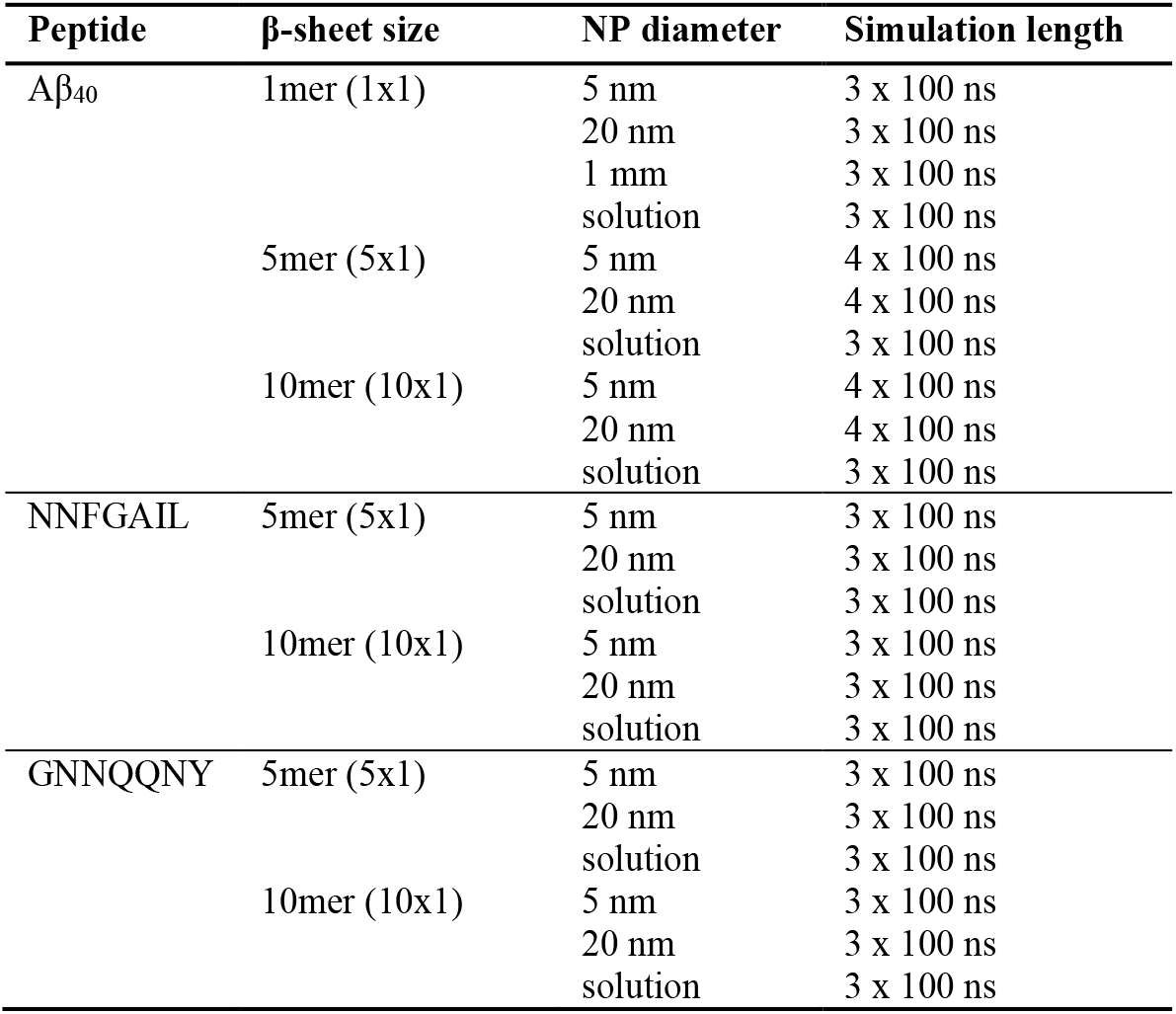
Overview of MD Simulations with Implicit NPs.

The conformational entropy of the Aβ_40_ 1mer (C_α_ atoms) for the curvature model and in solution was determined based on the Schlitter formula using the GROMACS tools g_covar and g_anaeig.^81^ Average absolute forces on the peptide N-termini and curvature potential energies were extracted from the simulations. Differences in absolute forces and potential energies between the NP-5 and NP-20 (difference = value for 5 nm – value for 20 nm) during the last 90 ns of simulation time (10-100 ns) were determined. Root-mean-square fluctuations (RMSF, g_rmsf) of the peptide residues were calculated in comparison to their respecting starting configurations for both the whole simulation trajectories and for the last 10 ns of simulation time. Relative RMSF values were corrected for the overall movement of the peptides by setting the RSMF minimum to 0 in each sequence. This enabled a comparison of the relative changes in structures between the simulation in solution and those bound with an NP-5 and NP-20. The β-sheet (and β-bridge) secondary structure content of the peptide aggregates was analyzed using the DSSP tool (Define Secondary Structure of Proteins, do_dssp) for the last 10 ns of simulation time.^82,83^ Averaged data of the repetitions are presented.

## ASSOCIATED CONTENT

### Supporting Information

Supporting methods (characterization of nanoparticles by dynamic light scattering, zeta potential measurements and UV-vis absorption spectroscopy; stability of gold nanoparticles in buffer solutions; prediction of peptide aggregation propensities; scanning electron microscopy), supporting results (characterization of nanoparticles by dynamic light scattering, zeta potential measurements and UV-vis absorption spectroscopy; stability of NP in buffer solutions; prediction of peptide aggregation propensities; Thioflavin (ThT) fluorescence assays; DLS, AFM and SEM of amyloid fibrils; Transmission electron microscopy (TEM) images; supporting MD simulation results, including Figures S1−S11 and Tables S1−S8), supporting references.

This material is available free of charge via the Internet at http://pubs.acs.org.

## AUTHOR INFORMATION

### Author Contributions

The study was designed by TJ, LLM, HRJ and BA. Experiments were performed by TJ (ThT fluorescence assays, DLS, UV-vis), JA (ThT fluorescence assays), JP (DLS, AFM, SEM) and MK (TEM). Nanoparticles were synthesized by CE (AuNP-mix). Experimental data were analyzed by TJ, JA and JP. Computer simulations were performed and analyzed by TJ and HRJ. The results were discussed and interpreted by TJ, JA, CE, DH, LLM, HRJ and BA. The manuscript was written by TJ and advanced by all authors.

### Notes

The authors declare no competing financial interest.

## ACKNOWLEDGMENT

The authors greatly acknowledge Dr. Jan Griebel (Leipzig, Germany) and Nadja Schönherr (Leipzig, Germany) for nanoparticle size and zeta-potential characterization, and Dr. Sven Rothemund (Leipzig, Germany) for peptide synthesis.

## ABBREVIATIONS

Aβ: amyloid beta
AFM: atomic force microscopy
AuNP: gold nanoparticle
DLS: dynamic light scattering
DMSO: dimethyl sulfoxide
DSSP: define secondary structure of proteins
GolP: Gold-Protein
HEPES: buffer (4-(2-hydroxyethyl)-1-piperazineethanesulfonic acid)
hIAPP: human islet amyloid polypeptide
MD: molecular dynamics
NP: nanoparticle
PBC: periodic boundary conditions
PDB: protein data bank
PME: Mesh Ewald method
RMSF: root-mean-square fluctuation
SEM: scanning electron microscopy
SPC: simple point charge
SUP35: protein (translation termination factor eRF3)
TEM: transmission electron microscopy
ThT: thioflavin T
TRIS: buffer (tris(hydroxymethyl)aminomethane)
UV-vis: ultraviolet–visible spectroscopy
ζ: zeta potential

