## Supporting Information for "Size Matters: A Mechanistic Model of Nanoparticle Curvature Effects on Amyloid Fibril Formation"

### 1 Supporting Methods

#### 1.1 Characterization of nanoparticles

***Dynamic light scattering (DLS) and zeta potential measurements.*** Measurements were performed on a Zetasizer Nano ZSP (Malvern Instruments, Malvern, UK) in polystyrene disposable cuvettes at 25°C. DLS was used to determine the relative hydrodynamic diameter (size) of the nanoparticles and was recorded at a wavelength of $\lambda$=633 nm and at a scattering angle of 173° for a duration of 120 s. The zeta potential ($\zeta$-potential) was measured with a zeta dip cell (ZEN1002) at a wavelength of $\lambda$=633 nm and a scattering angle of 173°. Dispersant was water (viscosity 0.8872 mPas, refractive index 1.330, and dielectric constant 78.5) and sample material were gold nanoparticles (AuNPs; refractive index 0.18, absorption 3.431).

***UV-vis absorption spectroscopy.*** Absorption spectra of the surface plasmon resonance of the nanoparticle solutions were recorded between 400 nm and 600 nm at a spectral resolution of at least 0.5 nm using an UV-2101PC UV-VIS spectrometer (Shimadzu, Kyoto, Japan).

#### 1.2 Stability of gold nanoparticles in buffer solutions

The stability of AuNP-mix (45 µg/mL), as a representative, was tested at different buffer concentrations (0.5 M, 0.25 M, 0.1 M, 0.05 M, 0.025 M) in phosphate, HEPES and TRIS buffer solutions (pH 7.4±0.05) to identify a buffer system that is suitable for the experiments with peptides, i.e. conditions under which the nanoparticles do not show major agglomeration. To measure the nanoparticle stability, UV-vis absorbance spectra were recorded in a transparent polystyrene well plate (Nunclon 96 Flat Bottom, Thermo Fisher Scientific, Waltham, MA, and Boettger, Bodenmais, Germany) between 350 nm and 700 nm at a spectral resolution of 2 nm at 30 °C after at least 10 minutes incubation time using an infinite 200 microplate reader (Tecan, Männedorf, Switzerland). HEPES buffer (0.1 M, pH 7.4) was chosen and the long-term stability of both AuNP-5 and AuNP-20 at 20 µg/mL was studied by measuring the UV-vis absorbance spectra after 30 minutes, 26 hours and 70 hours.

#### 1.3 Prediction of peptide aggregation propensities

Solubilities and aggregation propensities of the peptides were calculated using the CamSol[1,2] (www-vendruscolo.ch.cam.ac.uk/camsolmethod.html) and Aggrescan[3] (bioinf.uab.es/aggrescan/) methods. While CamSol predicts solubility in water, Aggrescan identifies aggregation-prone regions. Both methods are based on the physiochemical properties of each amino acid, its hydrophobicity, charge, or intrinsic aggregation propensity, in relation to their neighboring residues. The CamSol intrinsic solubility score results from the residues' solubilities, and the Aggrescan aggregation profiles (a4v) result from the amino-acid aggregation-propensity values (a3v) of each amino acid.

#### 1.4 Scanning electron microscopy (SEM)

Samples on silicon wafers were used as prepared for the atomic force microscopy (AFM) studies, detailed within the manuscript. SEM measurements were performed on an Ultra 55 microscope (Carl Zeiss Microscopy, Oberkochen, Germany) with an extra high tension (EHT) acceleration voltage of 2 kV, 3.6 mm free working distance and 20 µm aperture. SEM images were processed using SmartSEM 6.0 and Gwyddion 2.51 (http://gwyddion.net/).[4] Brightness and contrast were adjusted if appropriate.

### 2 Supporting Results

#### 2.1 Characterization of nanoparticles

All AuNPs were characterized using DLS (particle size, zeta potential) and UV-vis (surface plasmon resonance, SPR) measurements (see Table S1 and Figure S1). AuNP-5 and AuNP-20 were used for the peptide aggregation studies, while AuNP-mix served for initial experiments to identify a buffer system in which the AuNPs were stable.

**Table S1.** Overview of gold nanoparticles (AuNPs) and their UV-vis and DLS characterization.

| Name | UV-vis maximum / nm | Size obtained from UV-vis / nm | DLS size by number major species | DLS size by intensity all species >5% | zeta-potential / mV |
|---|---|---|---|---|---|
| **AuNP-5 [a]** | **<400 & 515** | **5 nm[5]** | **7 (±2) nm** | **11 (±4) nm, 97%** | **-41 (±10)** |
| **AuNP-20 [a]** | **524** | **20 nm[5]** | **17 (±4) nm** | **24 (±7) nm, 100%** | **-42 (±47)** |
| AuNP-mix [b] | 523 | 20 nm[5] | 1.3 (±0.3) nm | 32 (±12) nm, 66%<br>128 (±46) nm, 23%<br>1.5 (±0.3) nm, 9% | -25 (±14) |

[a] The size of the commercial nanoparticles (AuNP-5, AuNP-20) was confirmed by the manufacturer using electron microscopy. DLS is strongly influenced by the presence of small amounts of larger particles, and measures the hydrodynamic radius.

[b] AuNP-mix consist of mainly 1.5 nm and 32 nm large nanoparticles (two species). The signal at 128 nm is relatively low considering that larger particles result in a higher DLS signal even at low concentrations.

Higher SPR peak wavelengths in the UV-vis absorption spectra (Figure S1) indicate a bigger size of the nanoparticles.

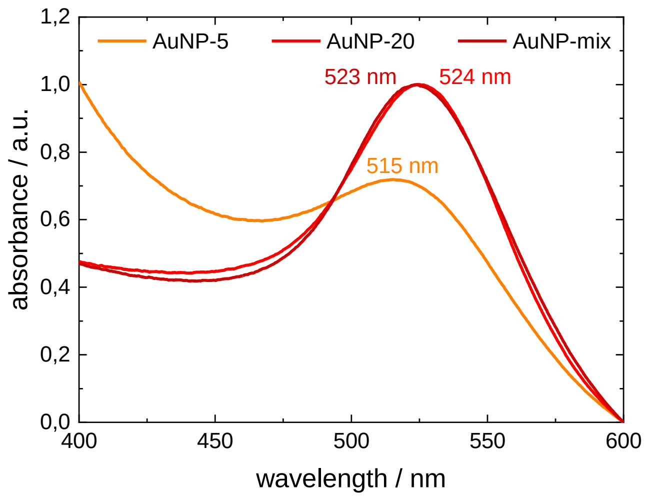


**Figure S1.** UV-vis absorption spectra of gold nanoparticles (AuNP-5, AuNP-20, AuNP-mix) in aqueous solution (highest absorbance each normalized to 1).

### 2.2 Stability of gold nanoparticles in buffer solutions

Colloidal nanoparticles without coating are unstable in solution and its surfaces are typically stabilized with carboxylic acids, alcohols or polymers.[6] The gold nanoparticles in this study (AuNP-5, AuNP-20, and AuNP-mix) were stabilized with a citrate-layer to prevent the nanoparticles from agglomeration by shielding the nanostructures using negatively charged surfaces. The addition of salts to nanoparticle solutions can induce their agglomeration if the charged surfaces are shielded.

The study of amyloid peptide aggregation should be performed under buffered conditions. To identify which buffer system and concentration can be used without any significant nanoparticle agglomeration, a screening of different conditions (phosphate, HEPES and TRIS buffer at concentrations between 0.025 M and 0.5 M) was tested using the AuNP-mix nanoparticles by UV-vis spectroscopy. The nanoparticle solutions were measured in water and various buffer solutions after at least ten minutes at 30°C. A red shift of the SPR spectra (i.e. a change to higher wavelengths) indicates an agglomeration and thus instability of the nanoparticles in the respective buffer. It was found that HEPES buffer can be used at 0.025/0.05/0.1 M concentration and phosphate buffer at 0.025 M concentration without nanoparticle agglomeration (see Figure S2). The AuNP-mix were not stable at any of the tested concentrations of TRIS buffer. HEPES buffer was chosen for the studies on amyloid peptide aggregation.

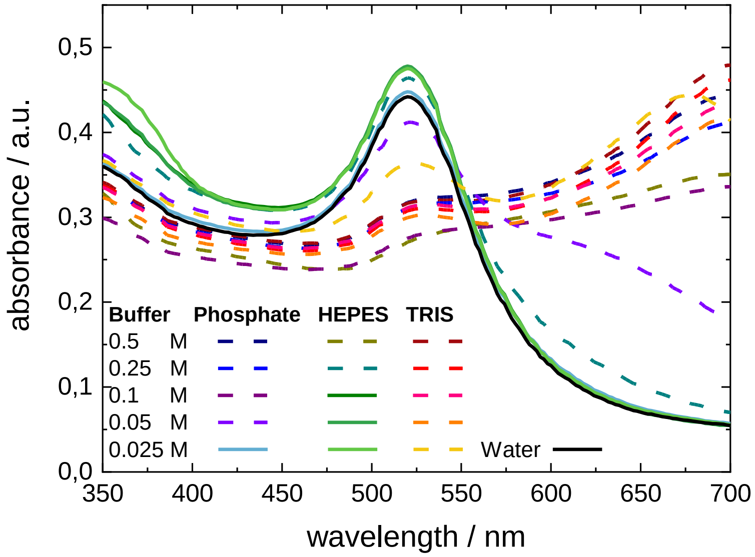


**Figure S2.** UV-vis absorption spectra of gold nanoparticles (AuNP-mix) in aqueous and buffer solutions (phosphate, HEPES and TRIS) at varying buffer concentrations (0.025-0.5 M). Spectra of stable nanoparticles are shown as lines, while agglomerated nanoparticles are shown as dashed lines.

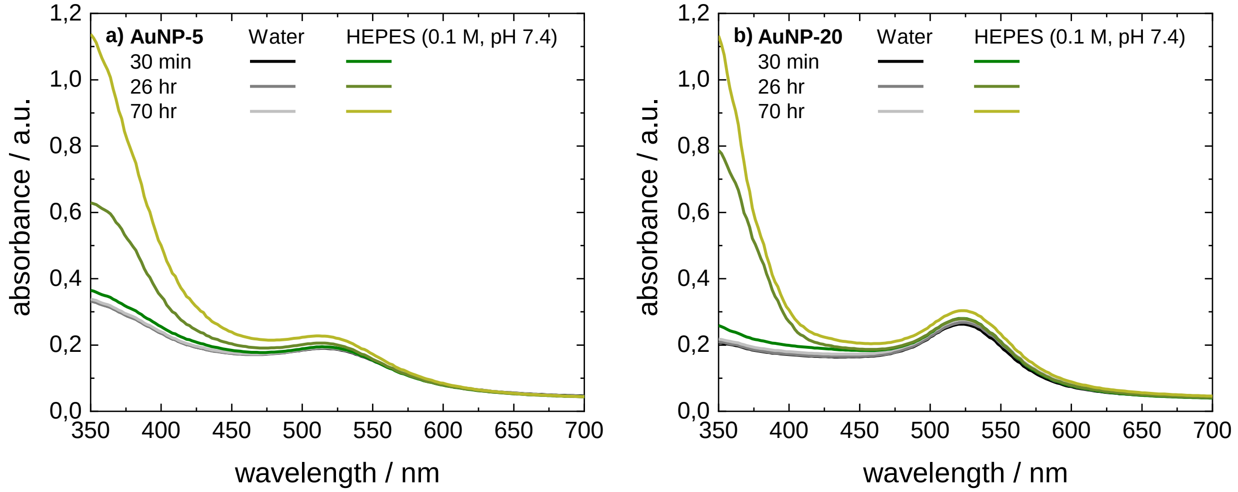


**Figure S3.** UV-vis absorption spectra of gold nanoparticles (a) AuNP-5 and (b) AuNP-20 in water and 0.1 M HEPES buffer solutions (pH 7.4). Both AuNP-5 and AuNP-20 remained stable upon addition of HEPES buffer for at least 70 hours. The absorption band below 400 nm only increased when HEPES buffer was present while the absorption of the nanoparticles remained unchanged.

Changes in solution conditions, such as pH value or salt concentration, can influence the aggregation kinetics.[7,8] To ensure a stable pH during the experiments, HEPES buffer was used (Table S2).

**Table S2.** pH values of water and 100 mM HEPES buffer solution before and after adding of AuNP-5 and AuNP-20.

| | **No NPs** | **AuNP-5** 20 µg/mL | **AuNP-20** 20 µg/mL |
|---|---|---|---|
| pH in water | 8.1 | 6.4 | 7.1 |
| pH in 100 mM HEPES | 7.4 | 7.4 | 7.4 |

### 2.3 Prediction of peptide aggregation propensities

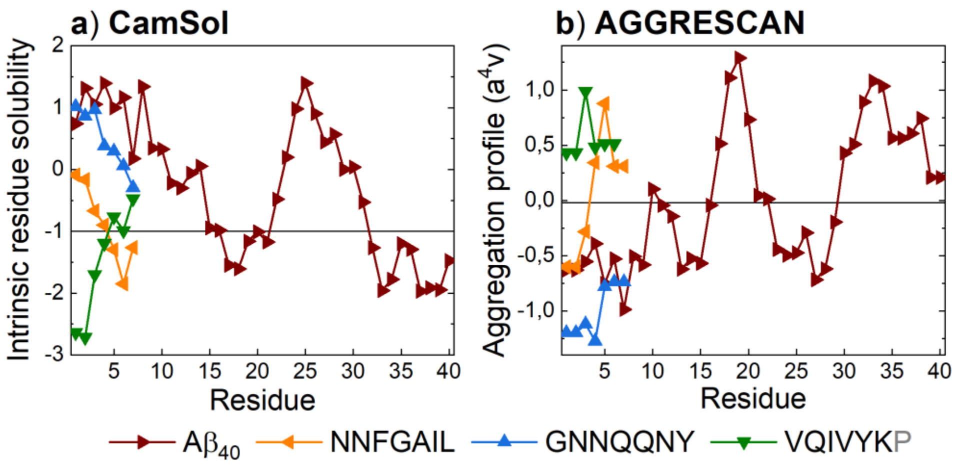


**Figure S4.** Prediction of (a) solubilities and (b) aggregation propensities of the studied peptides. (a) A low intrinsic residue solubility relates to a higher propensity to aggregate. Regions below -1 have a very low solubility. The CamSol method[1,2] only works for peptide sequences of at least 7 amino acids, which is why VQIVYKP was calculated (natural sequence). Overall, the peptides had the following intrinsic solubility scores: VQIVYKP (0.27) < $A\beta_{40}$ (0.66) < NNFGAIL (1.10) < GNNQQNY (1.89). (b) Higher values in the aggregation profiles[3], particularly above -0.02 are more prone to aggregation.

### 2.4 Thioflavin T (ThT) fluorescence

ThT fluorescence data for the peptides at 1 mg/mL are presented in Figure S5. An overview of half times of aggregation is summarized in Table S3.

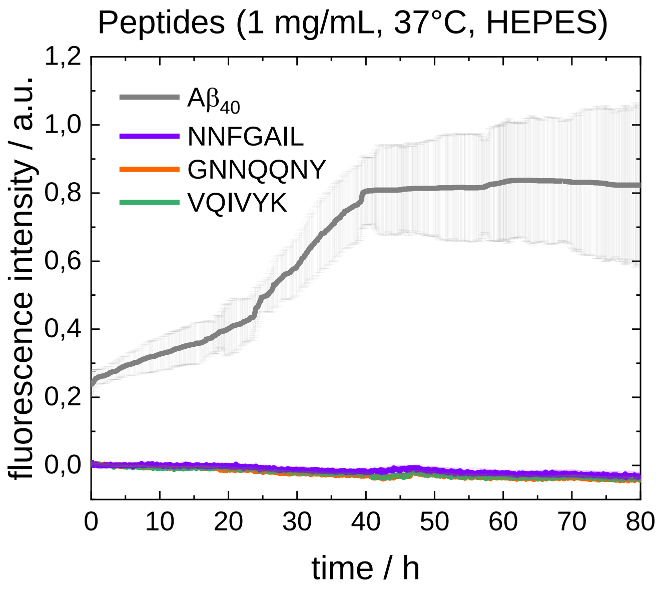


**Figure S5.** ThT fluorescence measurements to follow peptide aggregation in HEPES buffer (100 mM, pH 7.4). At 1 mg/mL, only $A\beta_{40}$ aggregated under the studied conditions within 80 h.

**Table S3.** Half times of aggregation ($t_{1/2}$, 50%) for $A\beta_{40}$, NNFGAIL, GNNQQNY and VQIVYK aggregation.

| **Sample** | **$t_{1/2}$ / h** **$A\beta_{40}$** 1 mg/mL, 37°C, HEPES | **$t_{1/2}$ / h** **$A\beta_{40}$** 1 mg/mL, 27°C, HEPES | **$t_{1/2}$ / h** **NNFGAIL** 3 mg/mL, 37°C, HEPES | **$t_{1/2}$ / h** **NNFGAIL** 3 mg/mL, 37°C, water | **$t_{1/2}$ / h** **GNNQQNY** 6 mg/mL, 37°C, HEPES | **$t_{1/2}$ / h** **VQIVYK** 10 mg/mL, 37°C, HEPES |
|---|---|---|---|---|---|---|
| **Control** | 29.0 | 46.5 | 0.16 | 7.5 | 11.0 | 10.5 |
| **AuNP-5 (20 µg/mL)** | 52.0 | > 80 | 0.11 | 1.8 | 25.5 | > 40 |
| **AuNP-5 (5 µg/mL))** | 32.5 | 53.5 | 0.06 | NA | 20.5 | > 40 |
| **AuNP-20 (20 µg/mL))** | 28.5 | 44.5 | 0 | 1.8 | 12.5 | 7.0 |

### 2.5 DLS, AFM and SEM of amyloid fibrils

DLS was used to follow the aggregation of $A\beta_{40}$ *in situ* without the presence of a fluorescence dye. When using this method, it needs to be noted that small amounts of any larger particles in a pure water or buffer solution scatter light very strongly (Figure S6). Thus, some measured intensities during the first minutes of measurement, when no larger fibrils were present, can be caused by the buffer itself. Further, this motivated to focus the analysis in the manuscript on the changes at larger hydrodynamic diameters.

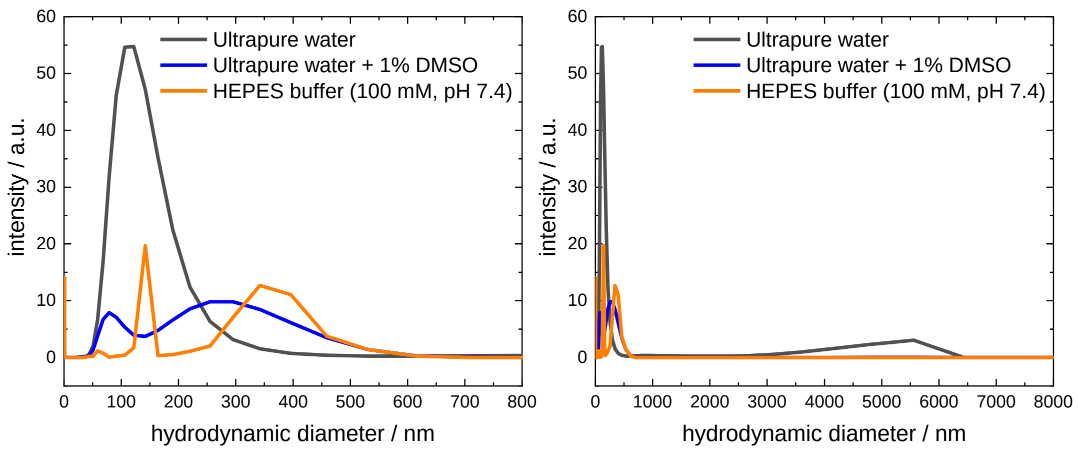


**Figure S6.** Dynamic light scattering (DLS) measurements of ultrapure water (grey), ultrapure water with 1% DMSO (v/v) (blue) and HEPES buffer (100 mM, pH 7.4) (orange) at 25°C as reference spectra.

Atomic force microscopy (AFM) and scanning electron microscopy (SEM) measurements were performed to follow $A\beta_{40}$ fibril growth (Table S4). No fibrils were visible at the start. After 8 days, fibrils were detected in all samples, with larger structures being present in the sample with AuNP-20. The AuNP-20 interconnected within the fibril structures (as previously shown for other peptides).[9]

**Table S4.** AFM and SEM images of Aβ40 peptide (1 mg/mL, 25°C) deposited onto silicon after incubation of peptide solutions in HEPES buffer without and with 20 µg/mL AuNPs (AuNP-5, AuNP-20) for eight days.

| | **AFM (warm scale) and SEM (gray scale) and images** |
|---|---|
| **Aβ40**<br>(1 mg/mL, 25°C)<br><br>**Control**<br><br>**8 days** | 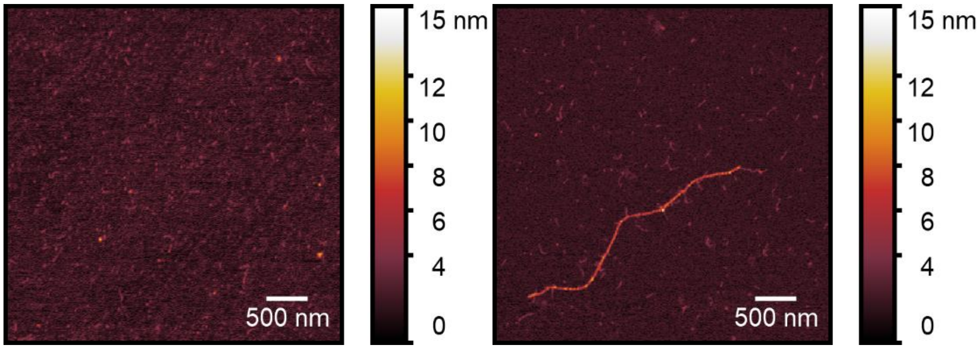<br> |
| **Aβ40**<br>(1 mg/mL, 25°C)<br><br>**AuNP-5**<br><br>**8 days** | 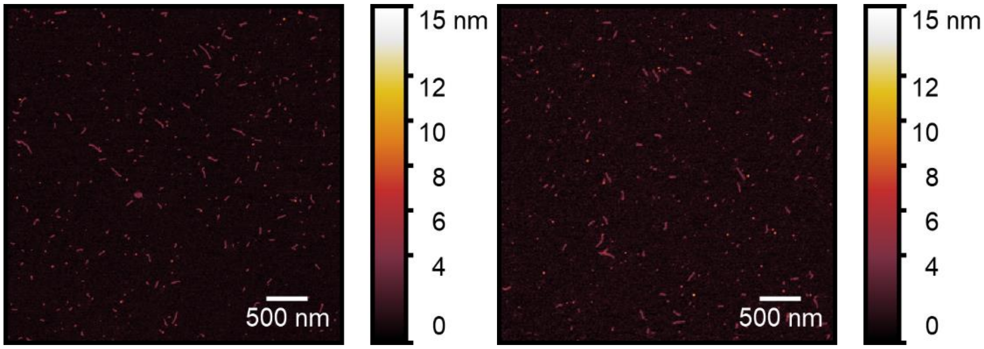<br> |
| **Aβ40**<br>(1 mg/mL, 25°C)<br><br>**AuNP-20**<br><br>**8 days** | 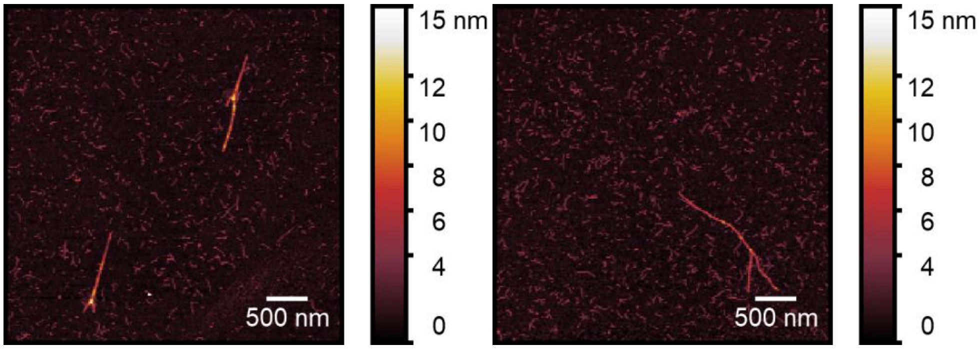<br> |

**A$\beta_{40}$**
(1 mg/mL, 25°C)

**Control**

**Start**

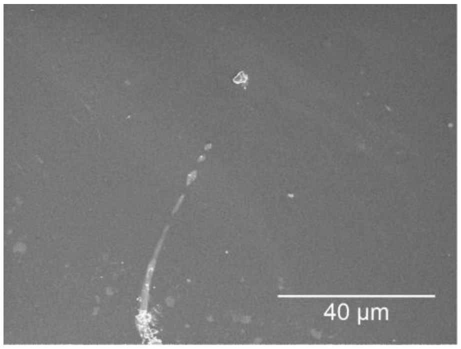


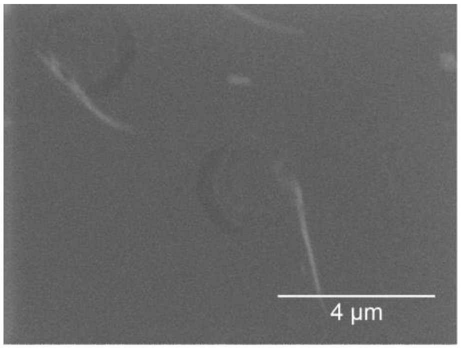


**A$\beta_{40}$**
(1 mg/mL, 25°C)

**AuNP-5**

**Start**

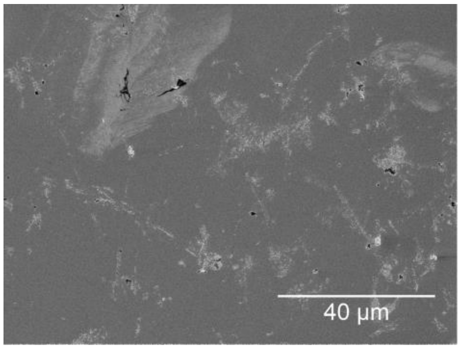


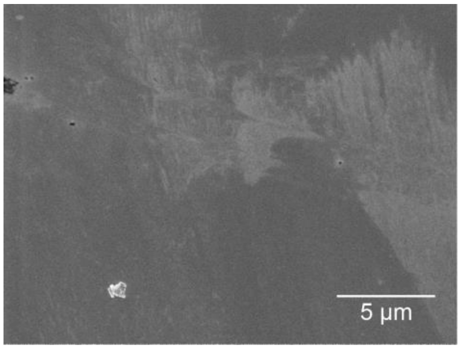


**A$\beta_{40}$**
(1 mg/mL, 25°C)

**AuNP-20**

**Start**

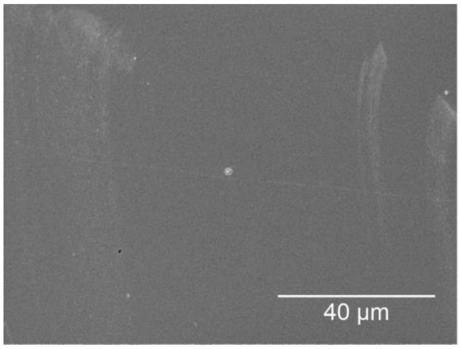


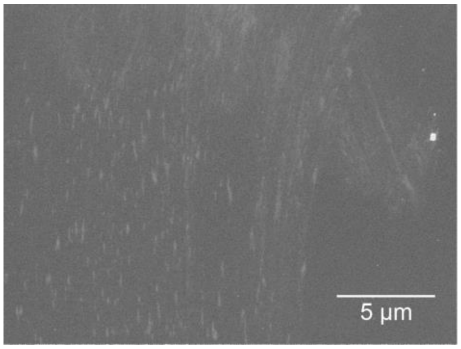


**A$\beta_{40}$**
(1 mg/mL, 25°C)

**Control**

**8 days**

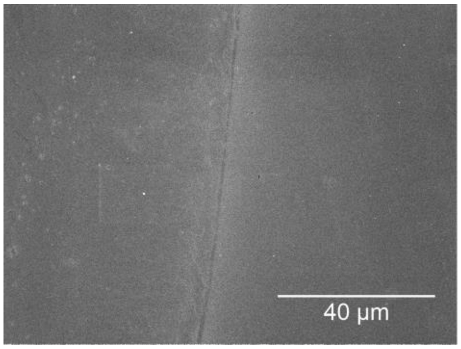


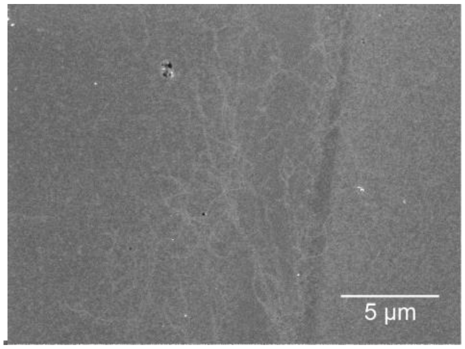

**A$\beta_{40}$**
(1 mg/mL, 25°C)

**AuNP-5**

**8 days**

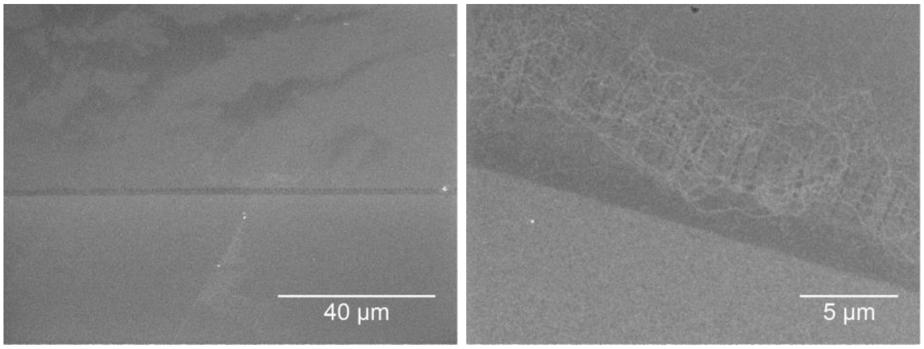


**A$\beta_{40}$**
(1 mg/mL, 25°C)

**AuNP-20**

**8 days**

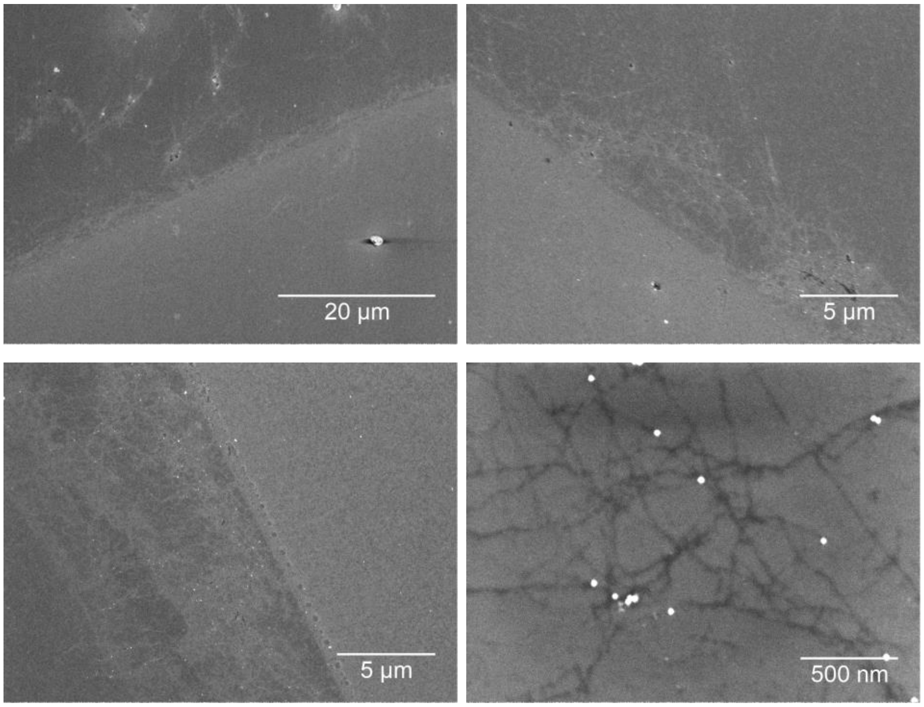


### 2.6 Transmission electron microscopy (TEM) images

The long-term formation of fibrils has been studied for A$\beta_{40}$, GNNQQNY and VQIVYK without and with AuNP-5 and AuNP-20 (20 µg/mL) present using electron microscopy (TEM) (Table S5). For NNFGAIL, fibril formation was previously reported.[9] Electron microscopy, as used here, will not reflect the kinetics but the morphology of formed fibrils. Samples were incubated for two months and air-dried on the substrate.

**Table S5.** TEM images of Aβ$_{40}$, GNNQQNY and VQIVYK peptide fibrils deposited onto silicon and air-dried after incubation of peptide solutions in HEPES buffer without and with 20 µg/mL NPs for two months (115 hours at 25/27°C shaken in ThT measurement, followed by 2 months in the fridge without shaking for slow fibril growth).

| | **Overview TEM image** | **Detailed TEM image** |
|---|---|---|
| **Aβ$_{40}$**<br>(1 mg/mL, 27°C)<br><br>**Control** | 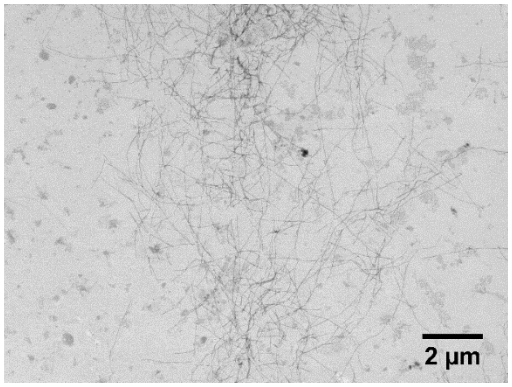 | 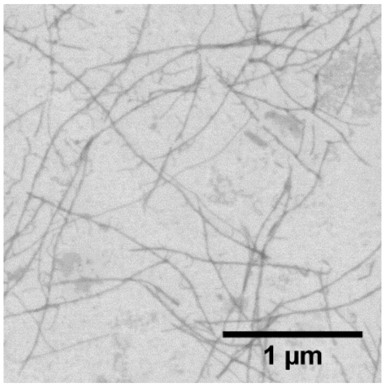 |
| **Aβ$_{40}$**<br>(1 mg/mL, 27°C)<br><br>**AuNP-5** | 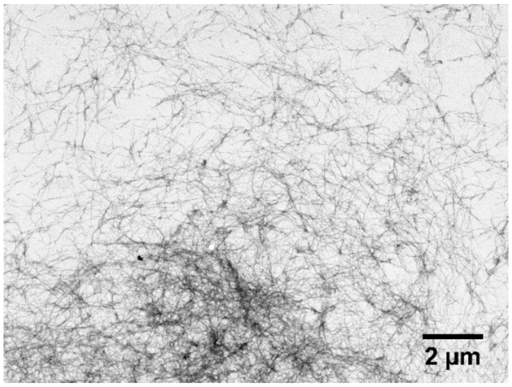 | 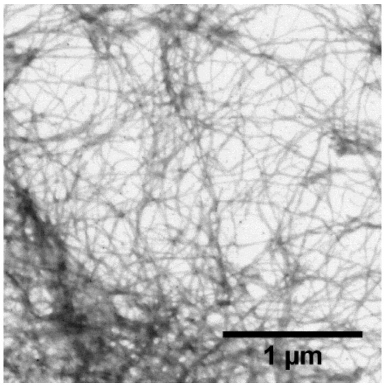 |
| **Aβ$_{40}$**<br>(1 mg/mL, 27°C)<br><br>**AuNP-20** | 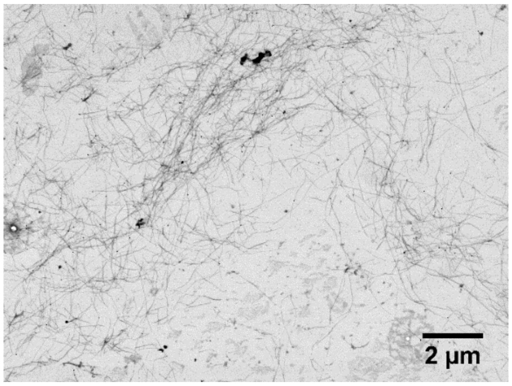 | 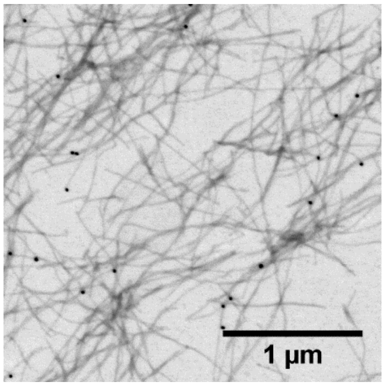 |

**GNNQQNY**
(4 mg/mL, 25°C)

**Control**

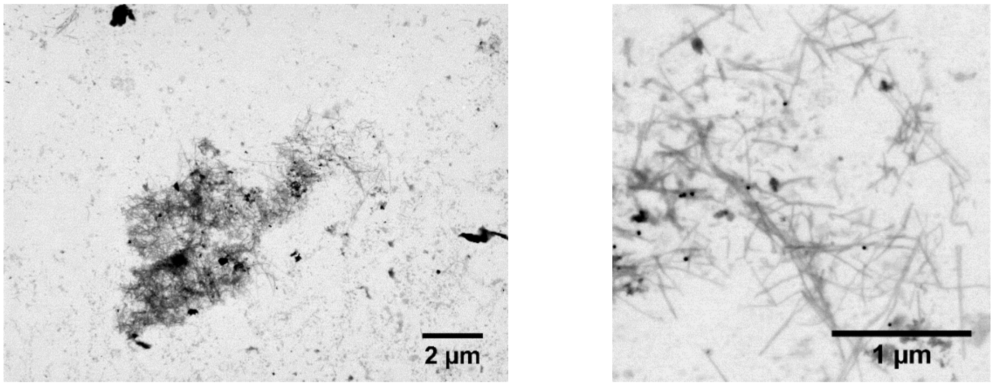


**GNNQQNY**
(4 mg/mL, 25°C)

**AuNP-5**

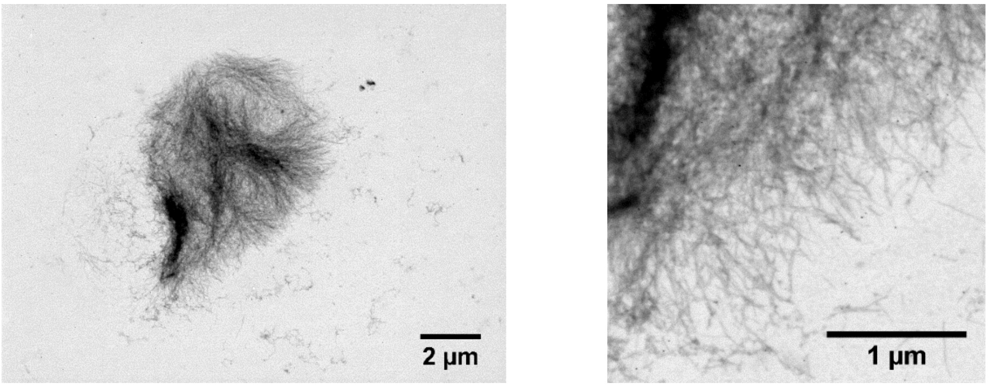


**GNNQQNY**
(4 mg/mL, 25°C)

**AuNP-20**

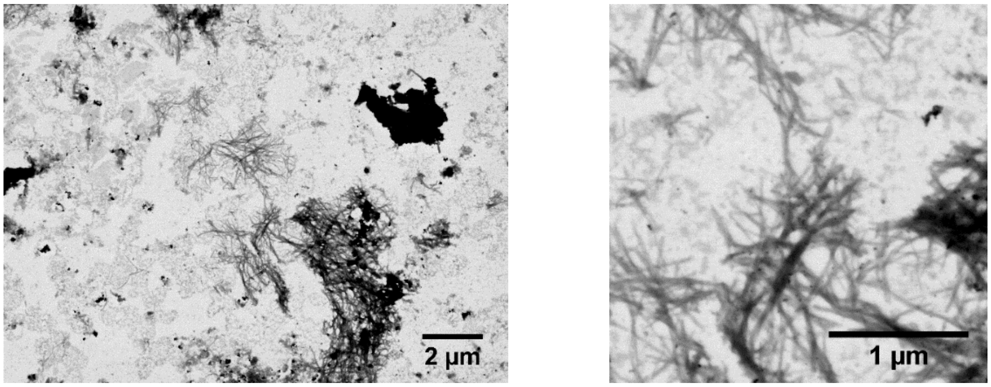


**VQIVYK**
(4 mg/mL, 25°C)

**Control**

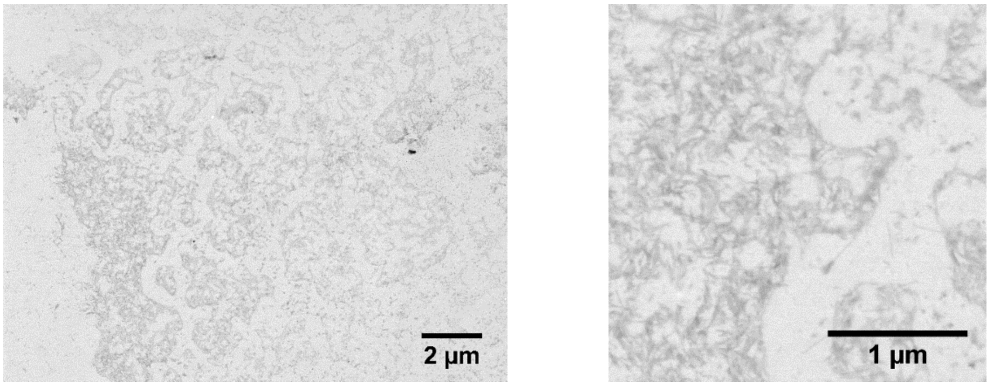

**VQIVYK**
(4 mg/mL, 25°C)

**AuNP-5**

**VQIVYK**
(4 mg/mL, 25°C)

**AuNP-20**

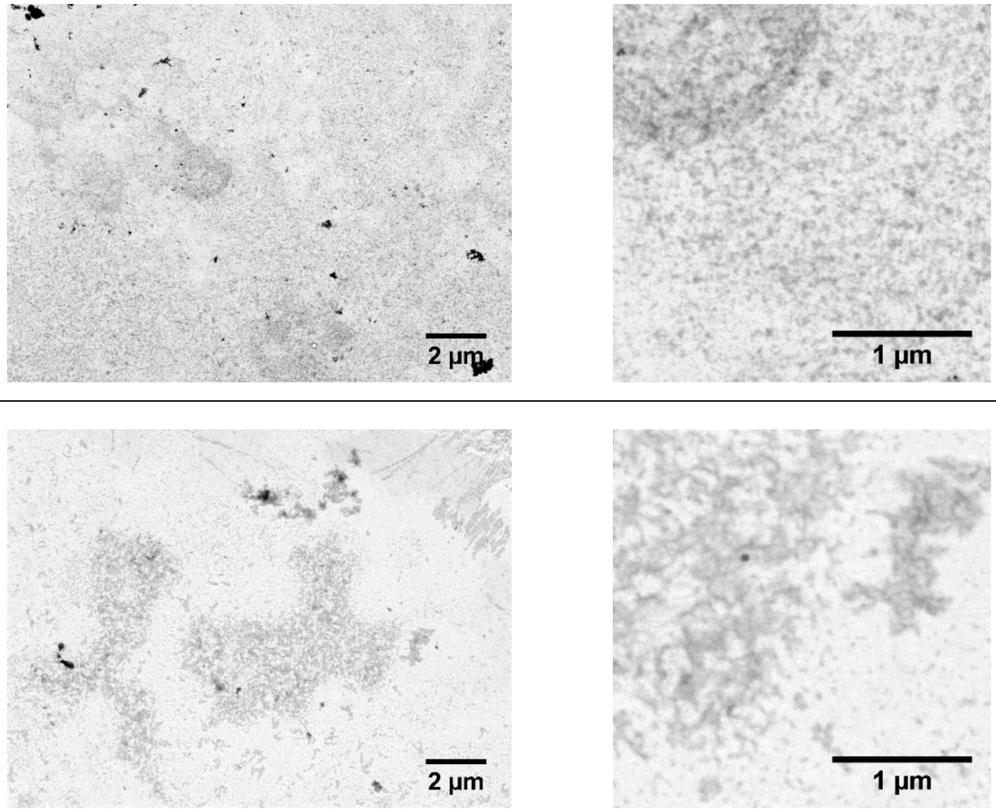


### 2.7 Supporting MD simulation results

The peptides A$\beta_{40}$, NNFGAIL and GNNQQNY were studied at citrate-coated gold surfaces using molecular dynamics simulations (MD) (see Figure S7). In initial simulations, several peptide monomers were placed in water without contact to the surface. Over time, the peptides adsorbed to the surface. These studies were previously reported for NNFGAIL, GNNQQNY and VQIVYK.[6,10] Bound peptide monomers from those simulations were used to study single peptides at the surface over time and to analyze their conformational flexibility compared to when the peptide monomers are free in solution.

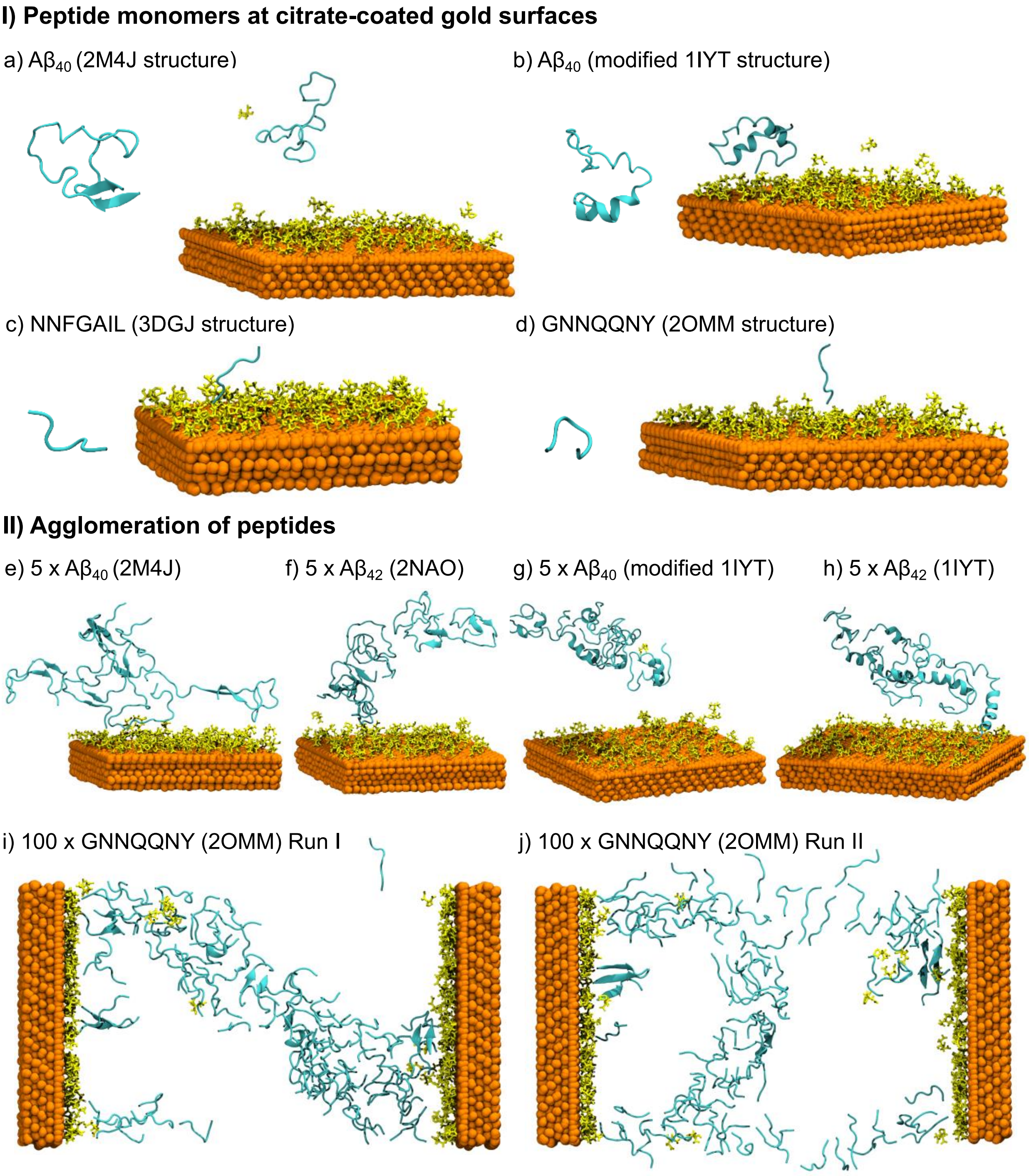


**Figure S7.** (I) MD simulation snapshots of peptide monomers of (a, b) $A\beta_{40}$, (c) NNFGAIL and (d) GNNQQNY at citrate-coated (shown in yellow) gold surfaces. One bound peptide (a-d) was simulated for 100 ns in multiple repetitions and compared with the peptide when simulated in water. Both (a) β-sheet and (b) α-helical starting structures were used for $A\beta_{40}$. (II) The further oligomer growth and peptide-surface interactions were studied for systems with multiple peptide atoms, both for (e, g) $A\beta_{40}$ and (f, h) $A\beta_{42}$ as well for (i - j) GNNQQNY. Data for GNNQQNY, NNFGAIL or VQIVYK have been published previously.[6,10]

The average secondary structure contents of the peptides during the last 10 ns of simulation time are summarized in Table S6 and Figure S8.

**Table S6.** The average secondary structure content of A$\beta_{40}$, A$\beta_{42}$, NNFGAIL and GNNQQNY peptide monomers and oligomeric structures during the last 10 ns of simulation time for the MD simulations with a citrate-coated gold-surface (@Au) and in solution.

| System | coil / % | β-sheet & β-bridge / % | bend / % | turn / % | α-helix / % | 3-helix /1 % |
|---|---|---|---|---|---|---|
| 1 x A$\beta_{40}$ (2M4J, β) @Au (9 rep.) | 46±2 | 11±2 | 34±3 | 9±2 | 0 | 0 |
| 5 x A$\beta_{40}$ (2M4J, β) @Au (1 rep.) | 54 | 15 | 26 | 6 | 0 | 0 |
| 1 x A$\beta_{40}$ (2M4J, β) in solution (3 rep.) | 50±4 | 9±4 | 36±3 | 5±1 | 0 | 0 |
| 1 x A$\beta_{40}$ (modified 1IYT, α) @Au (9 rep.) | 32±2 | 2±1 | 20±3 | 24±2 | 13±3 | 9±1 |
| 5 x A$\beta_{40}$ (modified 1IYT, α) @Au (1 rep.) | 34 | 1 | 17 | 23 | 16 | 8 |
| 1 x A$\beta_{40}$ (modified 1IYT, α) in solution (3 rep.) | 30±1 | 2±1 | 21±3 | 22±1 | 16±3 | 9±3 |
| 5 x A$\beta_{42}$ (2NAO, β) @Au (1 rep.) | 48 | 13 | 28 | 8 | 1 | 2 |
| 5 x A$\beta_{42}$ (1IYT, α) @Au (1 rep.) | 27 | 5 | 25 | 16 | 23 | 4 |
| 1 x NNFGAIL (3DGJ, β) @Au (9 rep.) | 81±4 | 0 | 18±3 | 1±0 | 0 | 0 |
| 1 x NNFGAIL (3DGJ, β) in solution (3 rep.) | 76±4 | 0 | 22±2 | 2±2 | 0 | 0 |
| 1 x GNNQQNY (2OMM, β) @Au (9 rep.) | 80±4 | 0 | 14±4 | 5±3 | 0 | 1±1 |
| 100 x GNNQQNY (2OMM, β) @Au (3 rep.) | 78±1 | 4±1 | 11±0 | 5±0 | 0 | 1±1 |
| 1 x GNNQQNY (2OMM, β) in solution (3 rep.) | 52±12 | 9±9 | 27±8 | 5±3 | 1±1 | 5±5 |

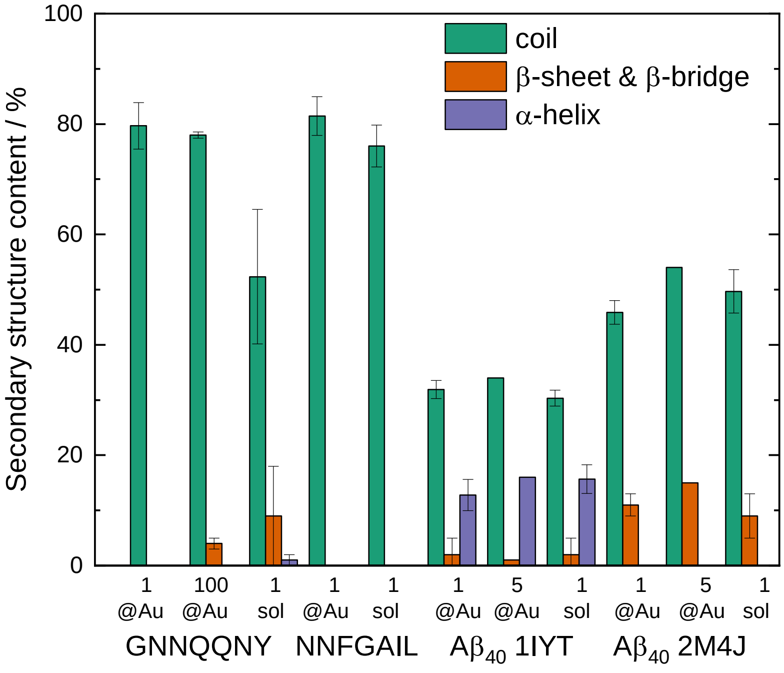


**Figure S8.** Visualization of the average secondary structure content of A$\beta_{40}$, NNFGAIL and GNNQQNY peptide monomers and oligomeric structures during the last 10 ns of simulation time for the MD simulations with a citrate-coated gold-surface (@Au) and in solution (sol).

The stabilities of linear parallel β-sheet rich aggregates consisting of five (5mer) and ten (10mer) peptide units of A$\beta_{40}$, NNFGAIL and GNNQQNY were studied in solution (Control) and attracted to an implicit curvature (5 nm and 20 nm diameter). In addition, one A$\beta_{40}$ monomer (1mer) was studied to compare to the simulations at planar surfaces. The peptide aggregate structures were constructed based on the fibril structure information obtained from the Protein Data Bank (see Methods section). While the aggregates consisting of five peptide monomers mimic early aggregates, the 10mers represent more mature prefibrillar species. Supporting simulation results for all simulations are shown in Figures S9 and S10.

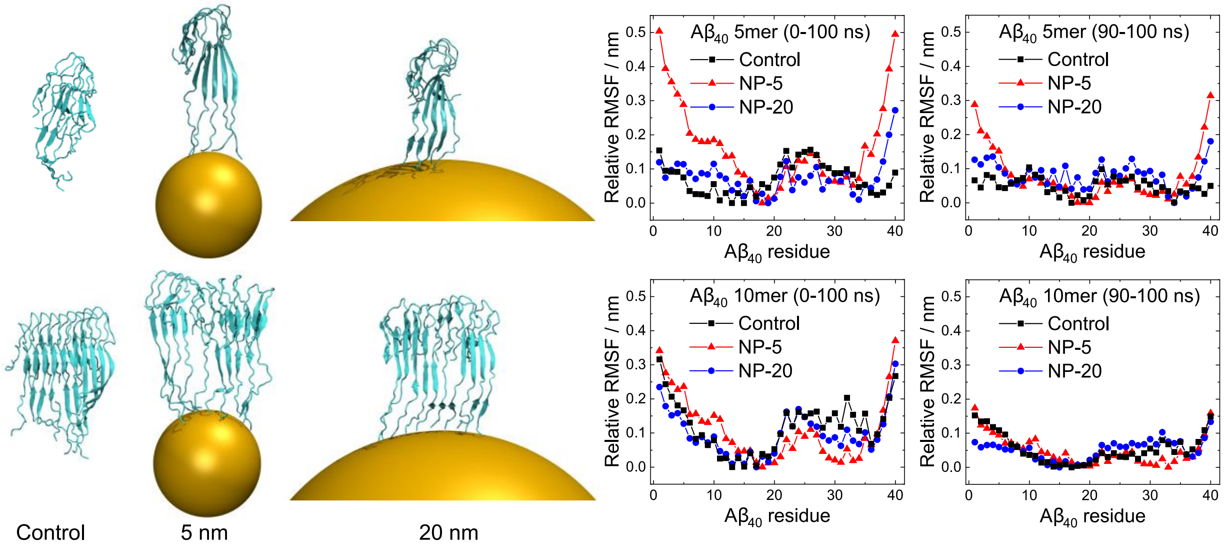


**Figure S9.** A$\beta_{40}$ aggregates (5mers and 10mers) were studied when bound to implicit nanoparticles of 5 nm and 20 nm diameter and when in solution (Control). The relative RMSF (root-mean-square fluctuation, relative = minimum for each condition set to 0 to correct for the overall movement) of the peptides over the simulation time of 100 ns and over the last 10 ns (90-100 ns) are shown. High RMSF values indicate large structural changes. The RMSF values shown during the last 10 ns are lower compared to those over the whole 100 ns of simulation time as the structures stabilized.

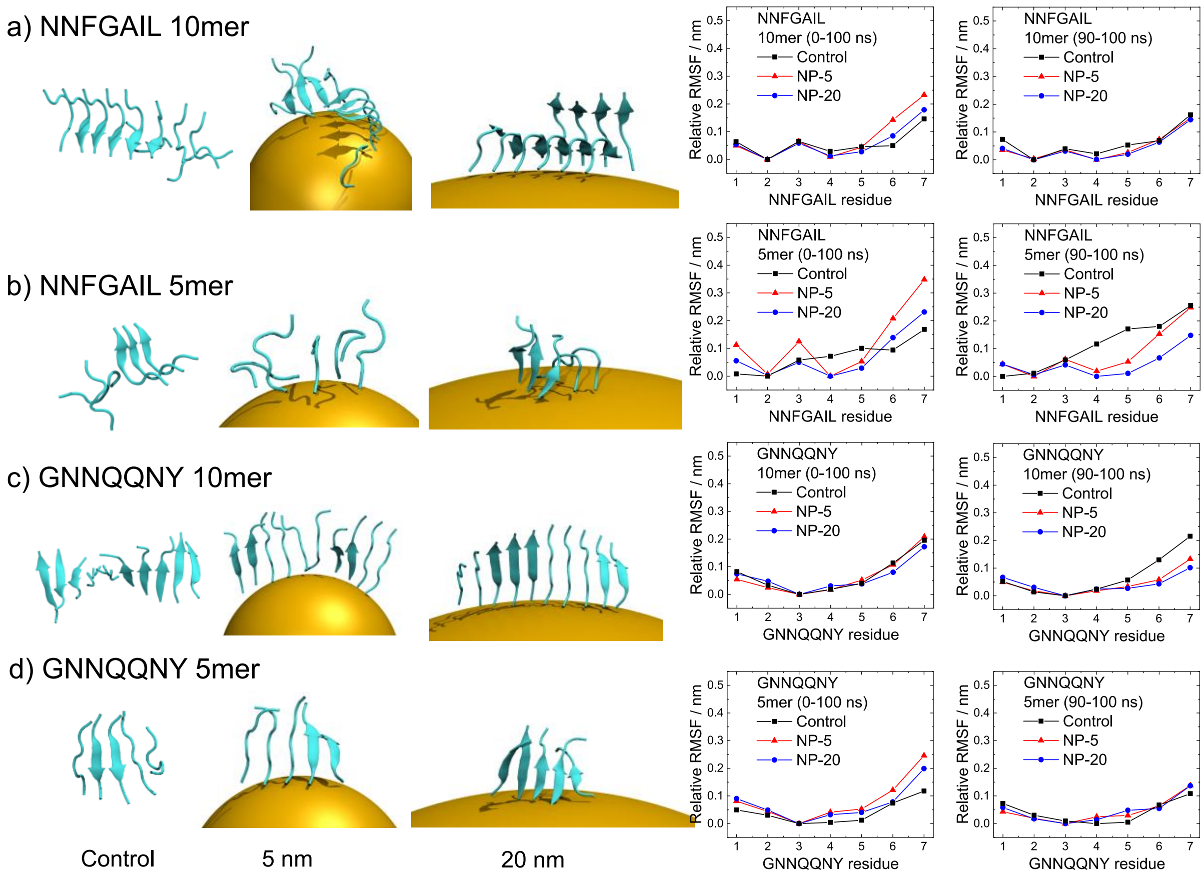


**Figure S10.** (a-b) NNFGAIL and (c-d) GNNQQNY aggregates, consisting of (a, c) 10 or (b, d) 5 peptide monomers, were studied when bound to implicit nanoparticles of 5 nm and 20 nm diameter and when in solution (Control). The relative RMSF (root-mean-square fluctuation, relative = minimum for each condition set to 0 to correct for the overall movement) of the peptides over the simulation time of 100 ns and over the last 10 ns (90-100 ns) are shown. Higher RMSF values indicate larger structural changes. The peptide C-terminus rearranged to a larger extent during the 100 ns simulation in the cases with NP curvature applied, while it was more stable during the last 10 ns where the peptides in solution show higher diffusion.

The average β-sheet contents of the peptides during the last 10 ns of simulation time are summarized in Table S7.

**Table S7.** The average β-sheet content of Aβ$_{40}$, NNFGAIL and GNNQQNY aggregates during the last 10 ns of simulation time for the MD simulations in solution (Control) and with an implicit curvature model (5 nm and 20 nm).

| System | Control β-sheet content / % | NP-5 β-sheet content / % | NP-20 β-sheet content / % |
|---|---|---|---|
| Aβ$_{40}$ 10mer aggregate | 27±4 | 28±2 | 24±1 |
| Aβ$_{40}$ 5mer aggregate | 20±1 | 28±3 | 22±4 |
| NNFGAIL 10mer aggregate | 16±1 | 16±7 | 15±4 |
| NNFGAIL 5mer aggregate | 9±2 | 1±1 | 11±7 |
| GNNQQNY 10mer aggregate | 48±1 | 51±2 | 51±3 |
| GNNQQNY 5mer aggregate | 45±3 | 48±5 | 52±1 |

The β-sheet content of the starting structures was as following: Aβ$_{40}$ 10mer: 19% (30% β-sheet + β-bridge), Aβ$_{40}$ 5mer: 14% (22%), NNFGAIL 10mer: 10% (28%), NNFGAIL 5mer: 0% (23%), GNNQQNY 10mer: 46% (46%), GNNQQNY 5mer: 41% (41%).

The NP curvature effect on the Aβ$_{40}$, NNFGAIL and GNNQQNY peptide aggregates (5mers, 10mers) was analyzed by quantifying the average absolute force on the N-termini and the NP potential energy over the last 90 ns (10-100 ns) of simulation time (see Table S8).

**Table S8.** The NP curvature effect on oligomers of the Aβ$_{40}$, NNFGAIL and GNNQQNY peptides was analyzed by quantifying the average absolute force on the N-termini and the NP potential energy over the last 90 ns (10-100 ns) of simulation time. Differences between the values at the smaller (NP-5) and larger (NP-20) NPs were calculated.

| System | Average absolute force on peptide N-termini $F$ / pN | | | NP potential energy $E_{pot}$ / kJ mol$^{-1}$ | | |
|---|---|---|---|---|---|---|
| | NP-5 | NP-20 | $F_5$ - $F_{20}$ | NP-5 | NP-20 | $E_{pot,5}$ - $E_{pot,20}$ |
| Aβ$_{40}$ 10mer | 22.97 ± 0.03 | 22.70 ± 0.03 | 0.28 ± 0.03 | 20.57 ± 0.06 | 20.79 ± 0.06 | -0.22 ± 0.06 |
| Aβ$_{40}$ 5mer | 21.35 ± 0.05 | 21.61 ± 0.05 | -0.26 ± 0.05 | 9.37 ± 0.04 | 9.54 ± 0.04 | -0.17 ± 0.04 |
| Aβ$_{40}$ 1mer | 20.08 ± 0.12 | 20.00 ± 0.12 | 0.08 ± 0.12 | 1.66 ± 0.02 | 1.65 ± 0.02 | 0.01 ± 0.02 |
| | NP-1000000: 20.56 ± 0.13; $F_{20}$-$F_{1000000}$: -0.56 ± 0.13 | | | NP-1000000: 1.78 ± 0.02; $E_{pot,20}$ - $E_{pot,1000000}$: -0.13 ± 0.02 | | |
| NNFGAIL 10mer | 20.09 ± 0.03 | 19.53 ± 0.03 | 0.56 ± 0.03 | 17.76 ± 0.06 | 16.69 ± 0.05 | 1.07 ± 0.06 |
| NNFGAIL 5mer | 21.00 ± 0.04 | 18.02 ± 0.04 | 2.98 ± 0.04 | 9.02 ± 0.04 | 7.12 ± 0.03 | 1.89 ± 0.04 |
| GNNQQNY 10mer | 20.43 ± 0.03 | 18.70 ± 0.03 | 1.73 ± 0.03 | 18.85 ± 0.06 | 15.95 ± 0.05 | 2.90 ± 0.06 |
| GNNQQNY 5mer | 20.41 ± 0.05 | 18.10 ± 0.05 | 2.31 ± 0.05 | 9.16 ± 0.04 | 7.25 ± 0.04 | 1.91 ± 0.04 |

The absolute values are largely curvature independent and determined by translational restraining and Brownian forces. Note that NP potential energy values will always be larger for larger fibrils and is force constant dependent. So value comparisons should only be done between NP sizes.

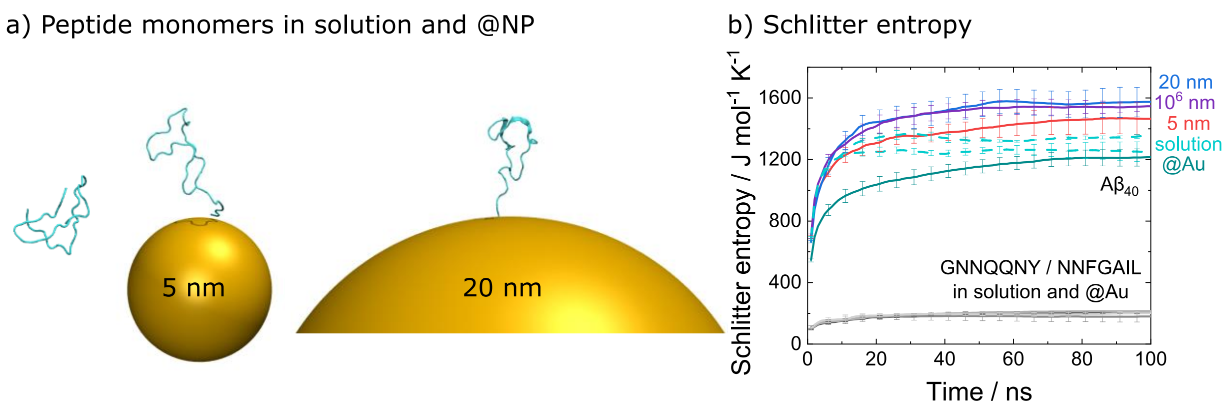


**Figure S11.** (a) A$\beta_{40}$ peptide monomers (1mers) were studied when bound to implicit nanoparticles of 5 nm and 20 nm diameter and when in solution. (b) The conformational entropy was analyzed using the Schlitter method (C$\alpha$ atoms for NPs of diameters 5 nm, 20 nm and 1,000,000 nm) and compared to the simulations of peptide monomers on planar citrate-coated gold surfaces (@Au) (see Figure 5 in manuscript).

(7) Haas, J.; Vöhrunger-Martinez, E.; Bögenhold, A.; Matthes, D.; Hensen, U.; Pelah, A.; Abel, B.; Grubmüller, H. Primary Steps of PH-Dependent Insulin Aggregation Kinetics Are

Governed by Conformational Flexibility. *ChemBioChem* **2009**, *10* (11), 1816–1822. https://doi.org/10.1002/cbic.200900266.

(8) Buell, A. K.; Galvagnion, C.; Gaspar, R.; Sparr, E.; Vendruscolo, M.; Knowles, T. P. J.; Linse, S.; Dobson, C. M. Solution Conditions Determine the Relative Importance of Nucleation and Growth Processes in α-Synuclein Aggregation. *Proc. Natl. Acad. Sci. U. S. A.* **2014**, *111* (21), 7671–7676. https://doi.org/10.1073/pnas.1315346111.

(9) Gladytz, A.; Wagner, M.; Häupl, T.; Elsner, C.; Abel, B. Structure-Making Effects of Metal Nanoparticles in Amyloid Peptide Fibrillation. *Part. Part. Syst. Charact.* **2015**, *32* (5), 573–582. https://doi.org/10.1002/ppsc.201400222.

(10) Gladytz, A.; Abel, B.; Risselada, H. J. Gold-Induced Fibril Growth: The Mechanism of Surface-Facilitated Amyloid Aggregation. *Angew. Chemie - Int. Ed.* **2016**, *55* (37), 11242–11246. https://doi.org/10.1002/anie.201605151.
